# SQuHIVLa: A novel assay for Specific Quantification of inducible HIV-1 reservoir by LAMP

**DOI:** 10.1101/2023.07.14.548928

**Authors:** Tanvir Hossain, Cynthia Lungu, Sten de Schrijver, Mamokoena Kuali, Shringar Rao, Ayanda Ngubane, Tsung Wai Kan, Robert-Jan Palstra, Paradise Madlala, Thumbi Ndung’u, Tokameh Mahmoudi

**Author notes:** ^8^ Authors contributed equally (Tanvir Hossain, Cynthia Lungu). Correspondence to: Tokameh Mahmoudi, Phone N.

## Abstract

Strategies toward HIV-1 cure aim to clear, inactivate, reduce or immunologically control the virus from a pool of latently infected cells such that combination antiretroviral therapy (cART) can be safely interrupted. In order to assess the impact of any putative curative interventions on the size and inducibility of the latent HIV-1 reservoir, robust and scalable assays are needed to precisely quantify the frequency of infected cells containing inducible replication competent HIV-1. Here, we present Specific Quantification of Inducible HIV-1 by LAMP (SQuHIVLa), a novel assay that leverages the high sensitivity and specificity of RT-LAMP, performed in a single reaction, to detect and quantify cells expressing Tat/Rev msRNA upon activation. Our LAMP primer/probe design exclusively detects subtype-specific HIV-1 Tat/Rev msRNA and exhibits high sensitivity, specificity, and reproducibility. Using SQuHIVLa we quantified the inducible viral reservoir in CD4+ T cells from a diverse group of people living with HIV-1 subtypes B and C on cART. SQuHIVLa presents a high throughput, scalable and specific HIV-1 reservoir quantification tool that is amenable to resource limited settings.

## Introduction

Combination antiretroviral therapy (cART) has changed human immunodeficiency virus type 1 (HIV-1) infection from a lethal to a chronic illness. However, despite over 40 years of research, there is currently no safe, scalable, curative therapy in sight. Furthermore, HIV-1 treatment disparities persist, with approximately 70% of all individuals with HIV-1 residing in sub-Saharan Africa^1^, and only 78% of them with access to treatment^2^. While standard clinical assays have shown that cART can effectively reduce circulating virus in the blood to undetectable levels, HIV-1 still persists in the form of an integrated provirus within a subset of immune cells, particularly resting memory CD4+ T cells^3, 4^ (the viral reservoir). If suppressive cART is interrupted, inducible replication-competent HIV-1 provirus (the inducible viral reservoir) can reactivate and exponentially replicate leading to detectable levels of viremia (viral rebound), within two weeks, in a majority of individuals^5^. Lifelong therapy is thus necessary to maintain viral suppression, and interventions beyond cART are urgently needed to achieve an HIV-1 cure.

Toward this aim, several strategies that target the inducible viral reservoir for clearance or control are being explored^6, 7^. To understand HIV-1 reservoir persistence, reliable biomarkers of reservoir size and dynamics are needed to evaluate the potential impact of interventions that aim to reduce the size of the inducible viral reservoir. Furthermore, robust, precise, and high throughput and scalable assays capable of measuring the frequency of infected cells containing inducible replication-competent HIV-1 are required^8^. However, the low frequency of these cells, estimated at between 1 and 1000 per million CD4+ T cells^9^ makes quantification extremely challenging, highlighting the need for innovative solutions to identify, target and quantify the viral reservoir in the quest for an HIV-1 cure.

Numerous techniques have been proposed, developed, or used to measure the HIV-1 reservoir^10 11, 12^. PCR-based assays offer a practical method to determine the frequency of cells containing total or integrated HIV-1 DNA ^13, 14^. However, these assays considerably overestimate the size of the viral reservoir. By targeting only one highly conserved proviral region, defective HIV-1 genomes that harbor large deletions or hypermutations in regions external to the assay amplicon are inclusively detected^15, 16^. Recent technological advances have addressed this issue by introducing assays that enable the detection of more than one conserved region of the HIV-1 genome. The intact proviral DNA assay (IPDA)^17^, for example, targets the packaging signal (PSI) and *env*-RRE regions using digital droplet (dd)PCR technology. Although successful in excluding the majority of defective viruses, conventional IPDA targets a very small sub-genomic region (2% of the HIV-1 genome), which can lead to an overestimation of the viral reservoir^18^. Adapted multiplex versions of IPDA have integrated additional assay targets for regions in HIV-1 *gag* and *pol* genes^19, 20^. A comparable technique to assess HIV-1 genome integrity, called Q4PCR, used a combination of a quadruplex qPCR and next-generation sequencing approach ^21^. While these assays have revealed novel insights into HIV-1 proviral landscape dynamics, some of the techniques are expensive, labor-intensive, and require sophisticated equipment, making them unsuitable for use in large-scale clinical trials, especially in resource-limited settings where the HIV-1 pandemic is more prevalent. In addition, these assays exhibit high assay fail rates due to HIV-1 sequence polymorphisms within the primer-probe binding regions^22, 23^ and near-full length genome sequencing methods may introduce a quantification bias^24^. Importantly, proviral DNA sequence intactness alone is not a direct measure of inducibility, which greatly depends on the site of viral integration in the host genome and therein the potential for viral transcription^16, 25–31^.

On the opposite end of the spectrum of reservoir quantitation, the culture-based, quantitative viral outgrowth assay (QVOA) is considered the gold standard technique to detect the outgrowth of replication-competent virus and therefore measures the functional viral reservoir ^32, 33^. However, QVOA underestimates the size of the viral reservoir due to suboptimal induction and/or inadequate propagation of all replication-competent viruses *in vitro*^16, 34^. Moreover, QVOA requires a high input of CD4+ T cells that necessitate large volume blood draws (>100 mL), is labor-intensive and time-consuming (14 to 21 days), limiting its use in routine clinical settings^35^.

Novel culture-based techniques have emerged as potential alternate assays to estimate the size of the inducible viral reservoir. These methods involve the activation of resting or total CD4+ T cells from virally suppressed individuals, followed by the direct measurement of HIV-1 RNA from cell extracts (cell-associated RNA, ca-RNA) ^36–39^ or cell culture supernatants (cell-free RNA, cf-RNA)^36, 37^. However, it should be noted that assays relying on quantification of cf-RNA may not accurately assess replication-competency as it only reflects the capacity to generate and release viral RNA^35, 36^. Inducible ca-RNA assays, on the other hand, involve the potent activation of CD4+ T cells by anti-CD3/CD28 antibodies^36, 37, 39^, phorbol 12-myristate 13-acetate (PMA) and ionomycin^38^, or latency reversing agents (LRAs)^36, 39, 40^, followed by an ultrasensitive PCR specific for a given viral transcript. Different RNA species including unspliced RNA (usRNA)^36, 40^, multiply spliced RNA (msRNA)^38, 39, 41^, mature RNA transcripts with poly-A tails^40, 42^, TAR RNAs^43^, and chimeric host-HIV-1 read-through transcripts^40, 41^ can be quantified by RT-qPCR or RT-ddPCR methods. Tat/Rev msRNA in particular has been found to be a meaningful indicator of viral replication following latency reversal^41, 44–47^ since Tat/Rev msRNA transcripts are generated after splicing of full-length viral transcripts. By detecting Tat/Rev msRNA therefore, the likelihood that proviruses with large internal deletions are measured is greatly reduced^16^. Inducible HIV-1 RNA assays can be conducted in a limiting dilution format, such as the Tat/Rev Limiting Dilution Assay (TILDA)^38, 48^, or at single-cell level by, for example, detection of HIV-1 usRNA and/or msRNA using fluorescent in situ hybridization assays coupled to flow cytometry (FISH-flow) which provide a deeper understanding of the viral reservoir molecular characteristics and phenotypic heterogeneity ^49–51^. However, these approaches still require high starting input (>50 mL of blood) and take 3 days to perform, which make FISH-flow assays less suitable for use in large scale cure intervention trials. Therefore, in most clinical studies viral RNA transcripts are preferably measured in bulk or in serial-diluted samples using quantitative real time or digital PCR-based methods^52^.

HIV-1 RNA quantification is typically performed using single round or semi-nested quantitative reverse transcription real-time PCR (RT-qPCR). Semi-nested RT-qPCR is especially useful for assessing samples with low copies of the target RNA and offers a wider dynamic range compared to single-round qPCR assays^41^. However, when HIV-1 RNA is quantified in limiting dilution format, e.g., in 96 wells or 384 wells, the cost of reagents considerably increases. Moreover, the instrument-intensive amplification procedure, long turnaround time, and high risk of cross-contamination from manual handling of pre-amplified product in semi-nested RT-qPCR protocols, limits assay application in large clinical trials, especially in resource-constrained settings^53^.

To address these technical limitations, reverse transcription loop-mediated isothermal amplification (RT-LAMP) could be used as a highly sensitive, less expensive and time-saving alternative to semi-nested RT-qPCR^54^. RT-LAMP uses two different polymerase enzymes to promote reverse transcription and isothermal amplification in a single reaction, which significantly reduces the risk of cross-contamination. RT-LAMP has been successfully used in multiple studies to detect low copies of viral RNA for SARS-CoV-2 diagnosis, demonstrating comparable or better sensitivity than conventional RT-qPCR^55–58^. Additionally, by targeting six to eight different DNA regions, RT-LAMP is more specific than semi-nested PCR, which targets three regions. Although RT-LAMP has the potential to be a less expensive and robust alternative to (semi-nested) RT-qPCR or RT-ddPCR^59^, it has thus far only been used for qualitative HIV-1 RNA detection in blood plasma^60–62^ and not yet in the context of HIV-1 reservoir quantification. Here, we present SQuHIVLa (Specific Quantification of Inducible HIV-1 reservoir by LAMP), a new HIV-1 reservoir assay that leverages RT-LAMP to detect HIV-1-infected cells that spontaneously or inducibly express Tat/Rev msRNA.

## Results

### A LAMP primer/probe set designed for exclusive detection of HIV-1 Tat/Rev msRNA

The successful design of a LAMP primer/probe set for specific LAMP detection of HIV-1 Tat/Rev msRNA, depended on a set of guidelines as depicted in Figure 1. The most critical criterion to ensure precise detection of Tat/Rev msRNA, excluding intron-containing Tat/Rev HIV-1 genomic DNA, was that the primer binding sites are exon-spanning. Using HIV-1 subtype B as reference, we designed primer/probe sets as depicted in the flowchart (Figure 1). We retrieved and aligned ten complete genome sequences of HIV-1 subtype B from different years (1983-2005) and various geographical locations to generate an *in silico*-spliced Tat/Rev HIV-1 DNA consensus sequence. Using the PrimerExplorer V5 software and the consensus Tat/Rev HIV-1 DNA FASTA file, we generated a list of preliminary sets of LAMP primers from which we selected primer sets where either the F2 binding region was within nucleotide position 125 to 240 or the B2 binding region was within nucleotide position 190 to 305, to ensure that the splice site (215^th^ nucleotide) was located within the forward loop (5’F2 to 3’F1) or backward loop (5’B2 to 3’B1). This criterion ensures that unstable loop formation resulting from primer binding to intron-containing Tat/Rev DNA, would inhibit isothermal amplification, thereby resulting in the exclusive detection of Tat/Rev msRNA (Figure 2A). Additionally, we generated two loop primers using the PrimerExplorer V5 software and aligned these and the other six LAMP primers to the ten HIV-1 genome sequences to assess primer-target sequence complementarity (Figure 2B). Given HIV-1’s high genetic diversity, it is impossible to ensure that the primer binding regions in patient-specific viral RNA are devoid of mutation hotspots. However, we strived to ensure that the mutation hotspots are located at positions such as the 5’ ends of F3, B3, F2, B2, and 3’ ends of F1c, B1c primer binding regions (Figure 2B). A set of primers with mismatches at these positions, whilst still meeting all the LAMP primer criteria such as melting temperature and distance between primer binding sites, would have the least impact on amplification (Methods). Furthermore, we converted the loop forward primer (LF) into a self-quenching probe to enable sequence-specific, real-time detection and quantification of LAMP amplicons (Methods).

**Figure 1.**
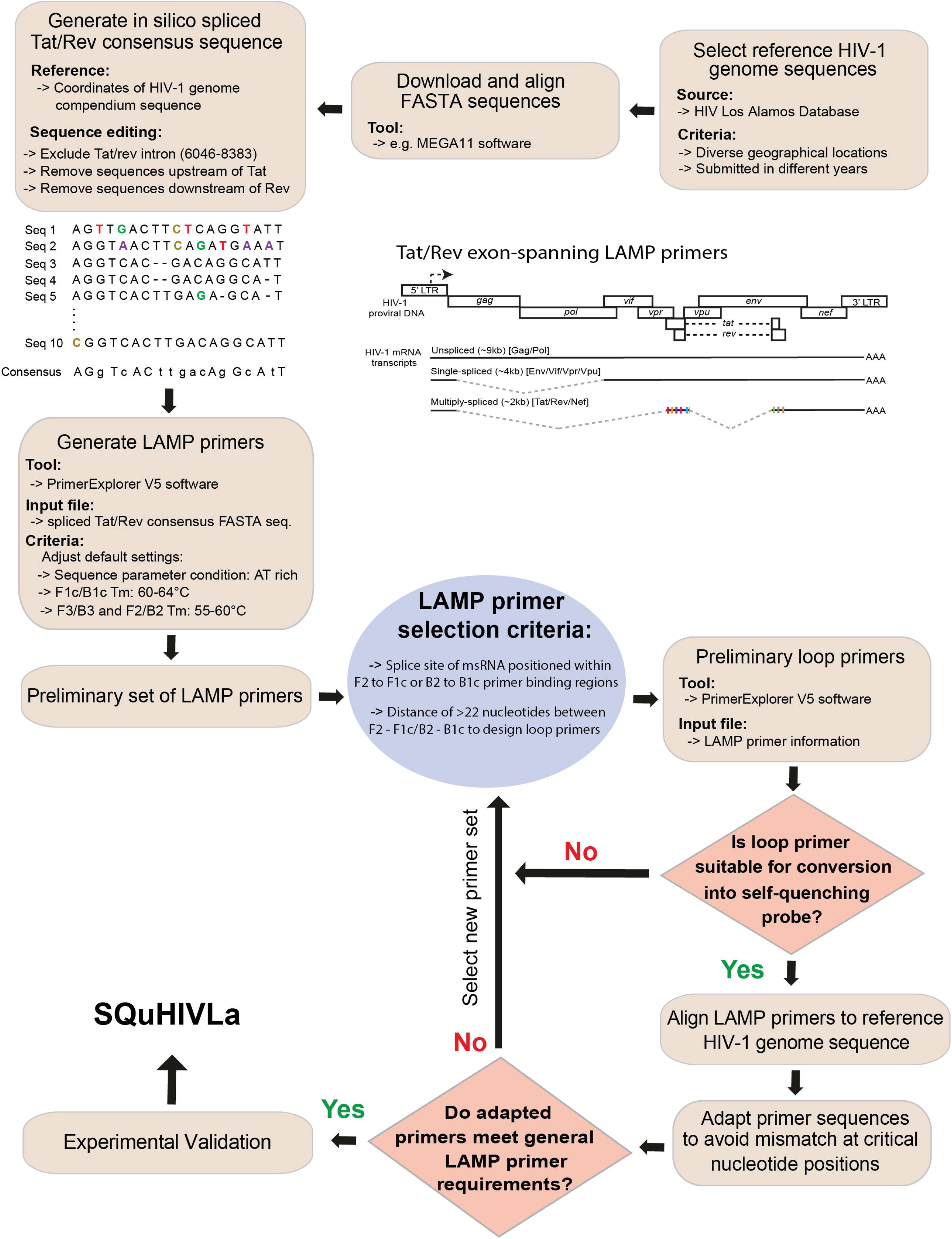
Flow chart of sequential steps in designing Tat/Rev msRNA LAMP primers/probe set.

**Figure 2.**
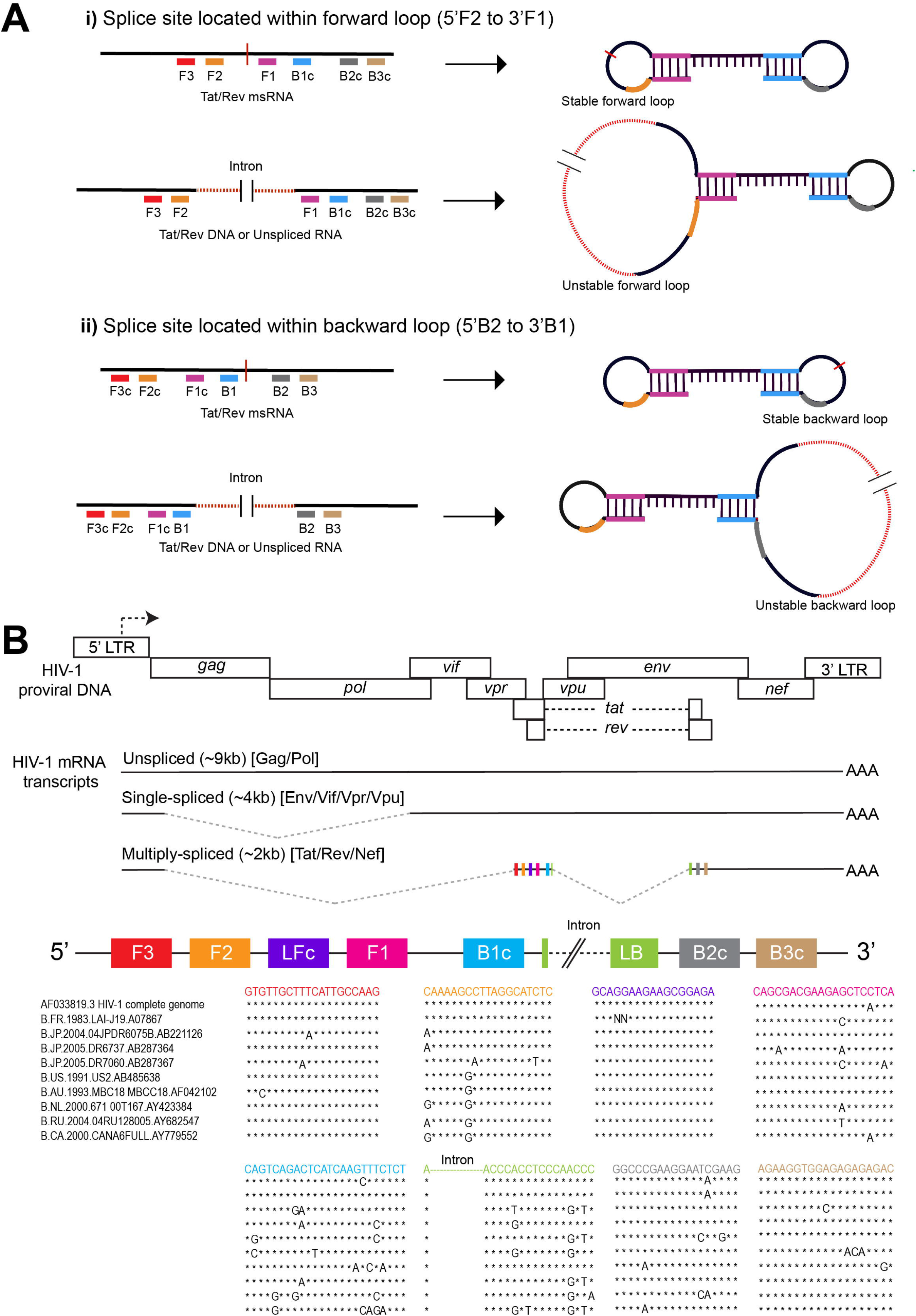
Schematic overview of SQuHIVLa and primer binding sites specific for HIV-1 subtype B. A) Schematic overview of the primer binding regions where the splice site is located within the forward or backward loop. Primers bound to msRNA results in a stable forward and backward loop formation. However, the presence of intron (in case of Tat/Rev DNA or usRNA transcript) leads to an unstable forward or backward loop formation, depending on the position of the intron within 5’F2 to 3’F1 or 5’B2 to 3’B1 respectively. B) Schematic overview of HIV-1 genome and transcripts produced during the HIV-1 replication cycle and alignment of the different LAMP primer binding regions with Ten HIV-1 subtype B viral sequences. The solid line represents the exons and the dotted lines represent the intron that is excised to generate mature HIV-1 mRNA transcripts. Exons of a multiply spliced RNA transcripts have colored rectangles that represent approximate LAMP primer binding sites. Similar to the alignment, different colors corresponds to different regions of the primers. After alignment, the "*" symbol is used to denote identical nucleotides; otherwise, the actual nucleotide symbol (A, T, C, or G) is used.

### Highly sensitive and specific detection of Tat/Rev msRNA by RT-LAMP

To assess the sensitivity of RT-LAMP in detecting Tat/Rev msRNA, we used serial dilutions of msRNA, *in vitro* transcribed from an intron-free Tat/Rev DNA template, driven by a T7 promoter (Figure 3A). As few as 50 copies of RNA were detected within 30.46 minutes (±7.21 minutes) using RT-LAMP in >97% of all reactions, with a Limit of Detection-95% at 31 copies (Figure 3B, 3D, 3E). At lower RNA copy numbers, RT-LAMP was less efficient in amplifying Tat/Rev msRNA shown by longer time to result and increased variability between reactions (Figure 3C). In experiments to assess the specificity of Bst 2.0 polymerase enzyme for Tat/Rev msRNA, we performed LAMP reactions with the required Bst 2.0 polymerase enzyme in the absence of the reverse transcriptase enzyme, RTx (Methods). Minimal and inconsistent amplification was observed at high copies of RNA (≥ 250 copies), suggesting some basal reverse transcriptase activity of the Bst 2.0 polymerase, which manufacturer validation experiments have also shown. However, the RT enzyme is necessary for robust and consistent amplification of RNA templates (Figure S1A and B).

**Figure 3:**
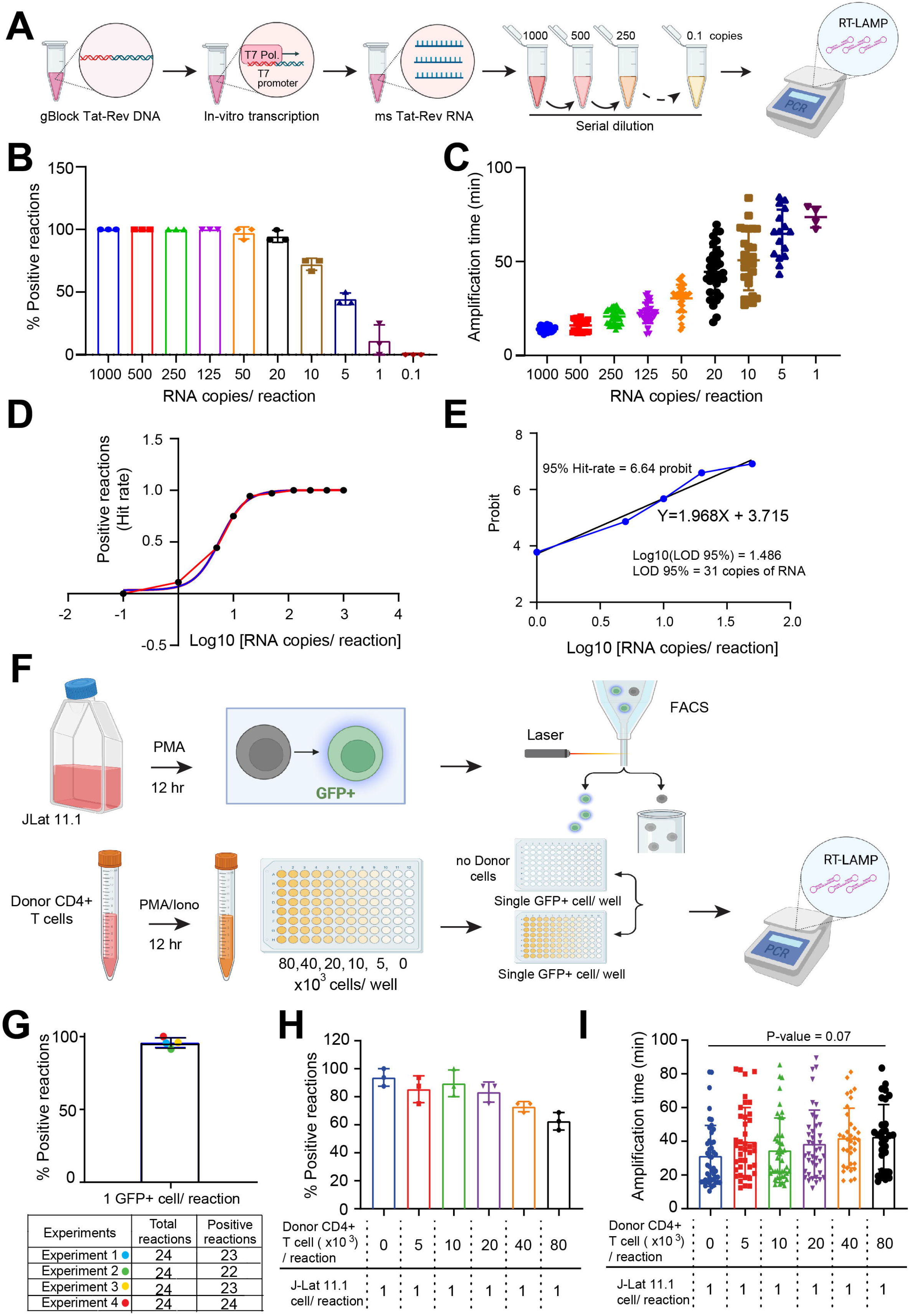
Sensitivity of RT-LAMP used in SQuHIVLa to detect msRNA. A) Experimental outline to generate synthetic msRNA using T7 polymerase mediated by *in vitro* transcription. Synthetic msRNA is serially diluted to achieve samples containing a range of target RNA copies used to perform RT-LAMP. B) Percentage of positive RT-LAMP reactions and C) amplification time required for a range of RNA copies are plotted. Data are presented as mean ± SD of three independent *in vitro* transcription experiments followed by serial dilution and RT-LAMP. D) A non-linear regression analysis is performed using the proportion of positive RT-LAMP reactions (Hit rate) (Y-axis) corresponding to Log10 values of RNA copies used as a template (X-axis). E) Probit analysis is performed to determine LOD-95% using probit values calculated from HIT-rate (Y-axis) corresponding to Log10 values of RNA copies (Y-axis). F) Experimental outline to sort single GFP+ J-Lat 11.1 cell in each well of 96 well PCR plate containing either no cells or different number of stimulated healthy donor CD4+ T cells to determine the sensitivity of RT-LAMP. G) Percentage of RT-LAMP positive reactions when a single GFP+ J-Lat 11.1 cell/reaction is probed in the absence of cellular background. Data are presented as mean ± SD of four independent experiments and each experiment is represented with different colored dots. H) Percentages of RT-LAMP positive reactions when a single GFP+ J-Lat 11.1 cell/reaction is probed in the presence of an increasing background of uninfected donor CD4+T cells and I) their corresponding RT-LAMP amplification time. Data are presented as mean ± SD of three independent experiments. Ordinary one-way ANOVA followed by Dunnett’s multiple comparison test are performed to analyze the variation of amplification times among different conditions and statistical significance is determined by p < 0.05.

Subsequently, to determine the sensitivity of RT-LAMP in detecting Tat/Rev msRNA directly from cells, we used J-Lat 11.1 cells, a Jurkat cell line harboring a latent but inducible full-length HIV-1 genome that expresses green fluorescence protein (GFP), as a product of msRNA transcripts (Figure S1C), upon activation. We sorted single GFP+ cells (Figure S1D) into RT-LAMP reaction plates in the presence or absence of PMA-stimulated uninfected donor CD4+ T cells (Figure 3F). In the absence of CD4+ T cell background, a single GFP+ cell was detected in 95.83% of all RT-LAMP reactions (Figure 3G), decreasing to 83.33% positivity in a background of 2x10^4^ cells, however the presence of cell background did not significantly impact the amplification time of RT-LAMP (mean amplification time ranging from 31.26 mins for no cell background to 42.63 mins in a background of 8x10^4^ cells) (Figure 3H and 3I).

Importantly, in another set of validation experiments we observed specific amplification of Tat/Rev msRNA and not intron-containing Tat/Rev genomic DNA with RT-LAMP (Figure 4A and 4B) while semi-nested PCR using non-exon spanning primers/probe detected both msRNA and genomic DNA (Figure S2). Minimal to no amplification was observed when RT-LAMP reactions were performed without the reverse transcriptase enzyme (Figure 4A, S3A) as well as in a range of validation samples including; GFP-J-Lat 11.1 cells that do not express msRNA (Figure 4A, S3B); DNase-treated J-Lat 11.1 RNA and RNase-treated J-Lat 11.1 DNA or using HIV-1 plasmid as template (Figure 4B). Overall, we demonstrate the rapid, sensitive and specific detection of cells expressing Tat/Rev msRNA by RT-LAMP and sought to use this assay for quantifying the inducible HIV-1 reservoir, hereafter referred to as SQuHIVLa (<u>S</u>pecific <u>Qu</u>antification of Inducible <u>HIV</u>-1 reservoir by <u>LA</u>MP).

**Figure 4:**
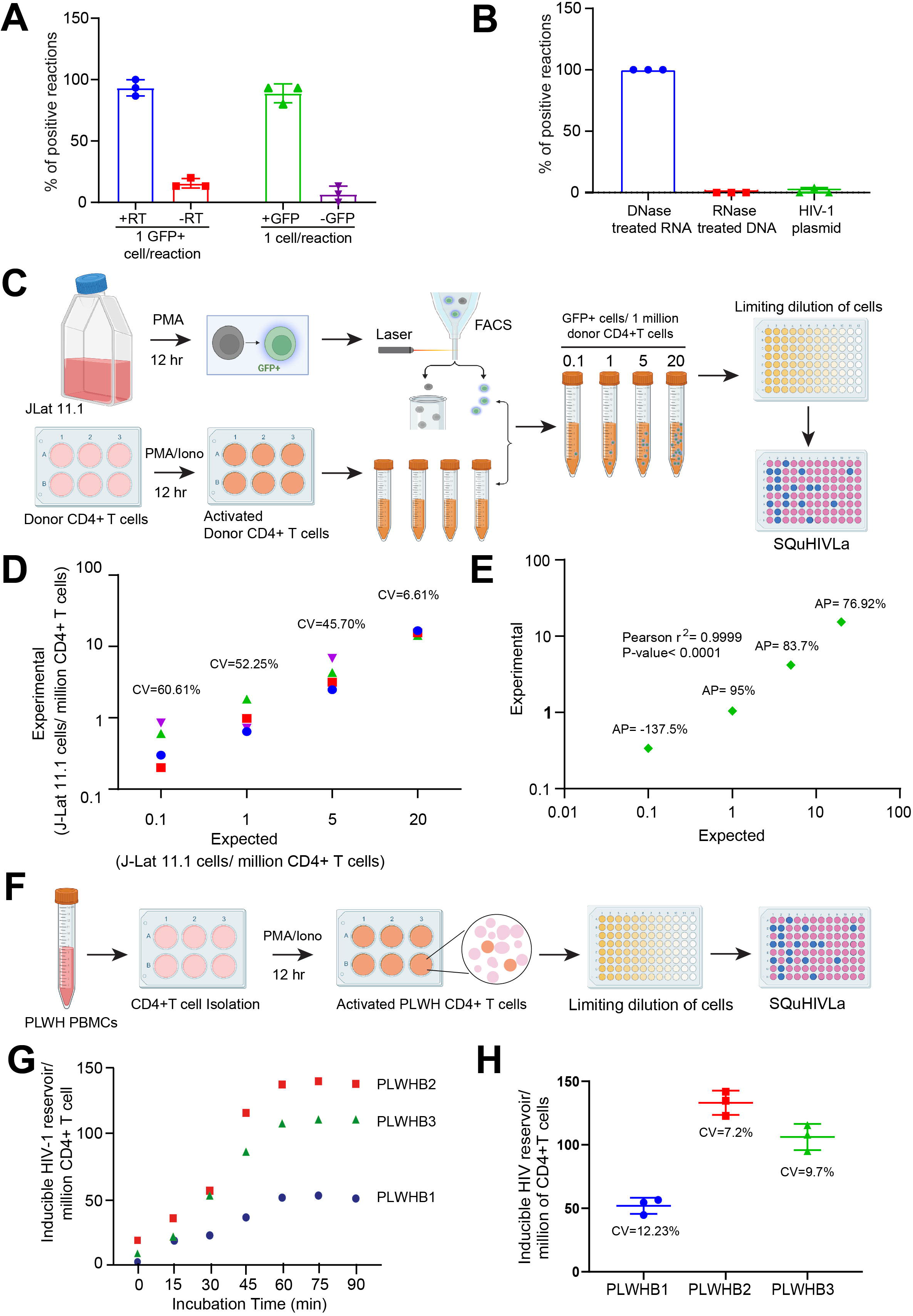
Quantification of inducible HIV-1 reservoir using SQuHIVLa. A) Percentages of RT-LAMP positive reactions when a single GFP+ J-Lat 11.1 cell/reaction was probed with or without reverse transcriptase enzyme included in the RT-LAMP reaction, and when either a single GFP+ or GFP-J-Lat 11.1 cell/reaction was probed. B) Percentages of positive reactions when a DNase-treated RNA and RNase-treated DNA sample isolated from PMA-stimulated J-Lat 11.1 cells, or when pNL4.3 E-R-plasmid were used as template for RT-LAMP. C) Experimental outline of preparing custom samples with an inducible HIV-1 reservoir of 0.1, 1, 5, 20 GFP+ cells/millions of CD4+ T cells using stimulated J-Lat 11.1 cells and uninfected donor CD4+ T cells. These custom samples were used to quantify the reservoir size using the SQuHIVLa assay. D) The inter assay coefficient of variation (CV) is determined from four independent experiments for all four samples representing different inducible reservoir size (0.1, 1, 5, 20 GFP+ cells/million CD4+ T cells). Four different colored-symbols represent data from four individual experiments. E) The Pearson correlation coefficient (r^2^) is determined between expected and experimental inducible reservoir size and statistical significance is determined by p < 0.05. The accuracy percentage (AP) was calculated with Excel using ABS function. F) A brief experimental outline of the application of SQuHIVLa to quantify inducible HIV-1 subtype B reservoir using PMA/ionomycin-activated CD4+ T cells in limited dilution format followed by maximum likelihood calculation. G) The inducible HIV-1 reservoir was quantified for three different people living with HIV-subtype B (PLWHB) using a SQuHIVLa protocol adapted to include an incubation at 45°C prior to RT-LAMP in order to determine the optimal incubation time for primary samples. Three different colored symbols represent data of three PLWHB. H) The inter assay CV is determined from three independent experiments for three PLWHB. Data are presented as mean ± SD and three different colored symbols represent three PLWHB.

### Accurate and reproducible quantification of the inducible viral reservoir using SQuHIVLa

In the next set of experiments, we sought to assess the reproducibility of SQuHIVLa in quantifying the inducible viral reservoir. To model HIV-1 reservoir samples, we generated a panel of four test samples comprising activated GFP+ J-Lat 11.1 cells (0.1, 1, 10 and 20 cells) sorted into a suspension of one million CD4+ T cells. Cells from each test sample were then distributed into RT-LAMP reaction plates in limiting dilution format, which allows quantification by maximum likelihood statistics (Figure4C, Methods). The observed and predicted viral reservoir size measurement were highly precise at frequencies greater than 20 GFP+ cells/million CD4+ T cells with a coefficient of variation (CV) of 6.61%. The CV increased to 60.61% at 0.1 cell GFP+ cells/million CD4+T cells (Figure 4D). Furthermore, the observed and predicted viral reservoir measurements correlated significantly with a good accuracy with mean accuracy percentage (AP) of 85% for reservoir size greater than 1 GFP+ cell/million CD4+ T cells (Figure 4E).

To further validate SQuHIVLa, we applied the assay to quantify the inducible viral reservoir in primary CD4+ T cells obtained from people with HIV-1 subtype B (PLWHB) on suppressive antiretroviral therapy. Following 12-hour PMA/ionomycin stimulation, which enhances transcriptional activation of inducible proviruses, whole CD4+ T cells were distributed in a limiting dilution format and Tat/Rev msRNA transcripts were amplified using RT-LAMP (Figure 4F). The frequency of cells expressing Tat/Rev msRNA was then determined by maximum likelihood estimation. We observed sub-optimal Tat/Rev msRNA amplification in primary CD4+ T cells, which was resolved through optimization of the sample incubation time at 45°C prior to isothermal amplification at 65°C; the viral reservoir size did not increase in three independent donors with incubation times exceeding one hour at 45°C, which was incorporated into the SQuHIVLa protocol when quantifying reservoir in primary CD4+ T cells (Figure 4G). The mean inter-assay CV, 9.58% (Range:7.2%-12.23%) was determined from three independent SQuHIVLa experiments using CD4+ T cells from three PLWHB demonstrating high reproducibility of SQuHIVLa in quantifying the inducible HIV-1 reservoir as depicted in Figure 4H.

### Adaptation of SQuHIVLa for non-B HIV-1 Subtypes

To determine the feasibility of using one primer-probe set for different HIV-1 subtypes, we aligned the subtype B Tat/Rev msRNA LAMP primer-probe sequences to genome sequences of HIV-1 subtypes C and A. A heatmap of the mismatch score, corresponding to the relative quantity of matching nucleotides for each primer binding region, revealed poor compatibility of the subtype B LAMP primer-probe sets for detection of subtypes C and A Tat/Rev msRNA (Figure S4). Several mismatches were present at sites critical for the LAMP reaction. Therefore, to specifically detect Tat/Rev msRNA from people with HIV-1 subtype C, globally the most prevalent subtype, we designed a new set of LAMP primers and probe (Figure 5A) following the aforementioned guidelines (Figure 1 and methods) and fulfilling the requirements to ensure specific amplification of subtype C Tat/Rev msRNA. The new primer-probe set demonstrated similar sensitivity in detecting *in-vitro* HIV-1 C Tat/Rev msRNA transcripts compared to the HIV-1 subtype B primer/probe set. In all experiments performed, 50 copies of Tat/Rev HIV-1 subtype C msRNA were detected in >97% reactions (Figure 5B) with a mean amplification time of 38.19 minutes (±7.47 minutes) (Figure 5C) and a LOD-95 % of 45 copies (Figure 5D, 5E), similar to that observed for Subtype B Tat/Rev msRNA.

**Figure 5:**
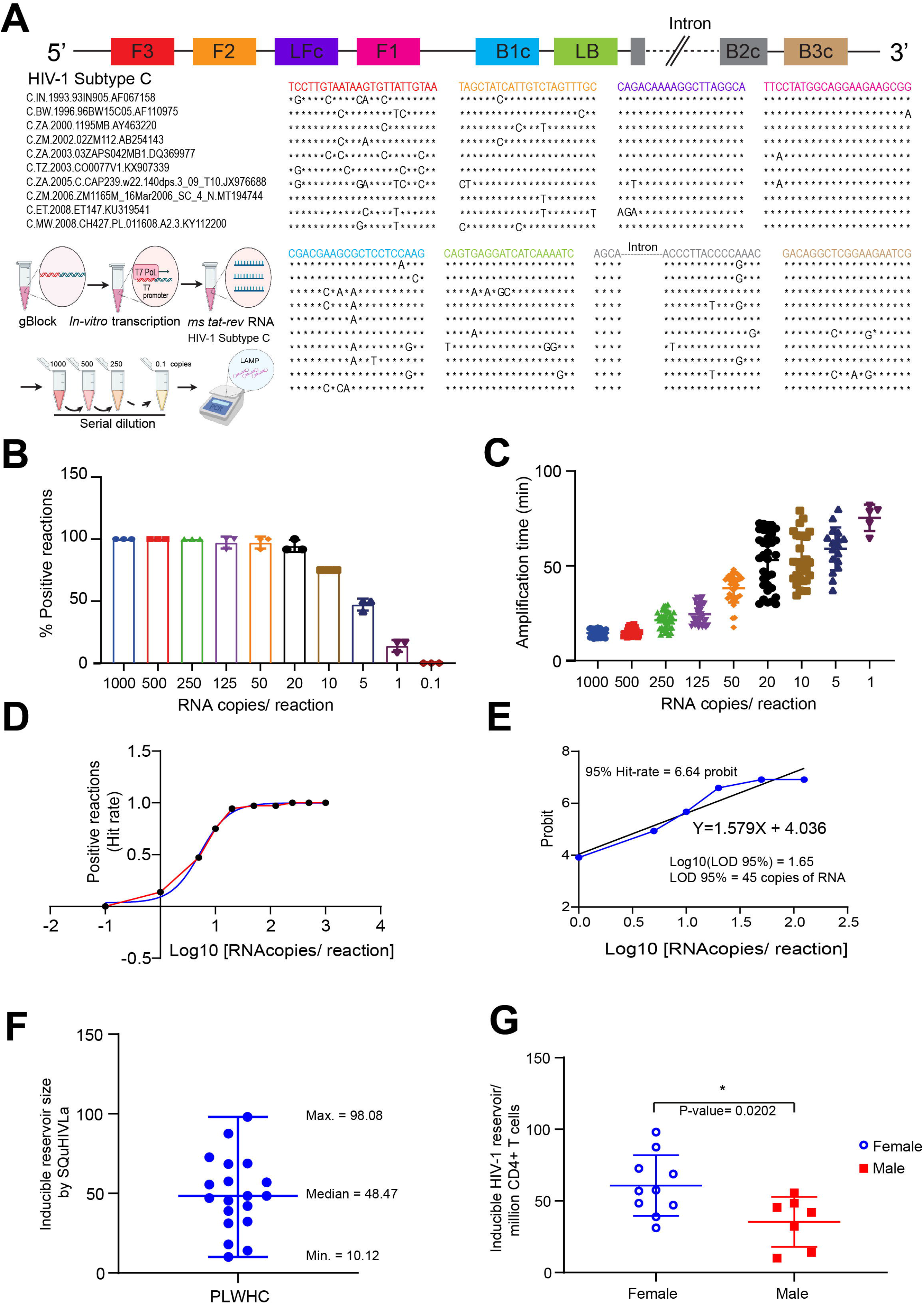
Design and application of SQuHIVLa to quantify the inducible reservoir of People with HIV-1 subtype C. B) Ten HIV-1 subtype C viral sequences are aligned with different subtype C specific LAMP primers. After alignment, the "*" symbol is used to denote identical nucleotides; otherwise, the actual nucleotide symbol (A, T, C, or G) is used. *In vitro* transcribed synthetic subtype C msRNA is serially diluted to achieve samples containing a range of target RNA copies used to perform RT-LAMP with subtype C specific primers and probe. B) Percentage of positive RT-LAMP reactions and C) amplification time required for different amounts of RNA copies are plotted. Data are presented as mean ± SD of three independent *in vitro* transcription experiments followed by serial dilution and RT-LAMP. D) A non-linear regression analysis is performed using proportion of positive RT-LAMP reactions (HIT rate) (Y-axis) corresponding to Log10 values of RNA copies used as a template (X-axis). E) Probit analysis is performed to determine LOD-95% using probit values calculated from the HIT-rate (Y-axis) corresponding to Log10 values of RNA copies (Y-axis). F) The inducible HIV-1 reservoir was quantified for N=19 people living with HIV-1 subtype C (PLWHC) presented as median with range. level G) The inducible HIV-1 reservoir was quantified for ten female (blue open circle) and seven male (red square) PLWHC using SQuHIVLa assay. Data are presented as mean ± SD. Two-tailed unpaired t-test is performed to analyze the variation of inducible HIV-1 reservoir between male and female PLWHC and statistical significance is determined by p < 0.05.

We then assessed the performance SQuHIVLa across a panel of clinical samples from 19 people living with HIV-1 Subtype C (PLWHC) on fully suppressive antiretroviral therapy (17 chronically infected and 2 elite controllers). Tat/Rev msRNA was detectable in all samples and the frequency of cells expressing Tat/Rev msRNA ranged from 10.12-98.08 cells per million CD4+ T cells (Figure 5F). Interestingly, in the chronically infected PLWHC, the inducible viral reservoir was notably higher in females (n=10) than in males (n=7) (P=0.0202) (Figure 5G). Additionally, viral reservoir quantification was also possible for the two HIV-1 Subtype C elite controllers (Table 1). Univariate correlation analyses did not reveal significant association between the frequency of cells expressing Tat/Rev msRNA and CD4+ T cell counts or pre-ART plasma HIV-1 RNA (Figure S5) Overall, these findings demonstrate that SQuHIVLa is highly adaptable and has great potential for application in studies involving people with HIV-1, with various clinical characteristics.

**Table 1.** Participant characteristics.

| Patient ID | Age<br>(years) | Sex<br>assigned<br>at birth | ART<br>regimen | Viral load at<br>sample<br>collection | CD4+ T cell<br>count<br>(cells/mm<sup>3</sup>) | Duration of<br>viral<br>suppression<br>(months) | CD4 nadir<br>(cells/mm<sup>3</sup>) | Pre-cART<br>plasma HIV-1<br>RNA (log10<br>copies/mL) | Reservoir<br>size<br>(SQuHIVLa) |
| --- | --- | --- | --- | --- | --- | --- | --- | --- | --- |
| HPP 37 | 25 | F | ODIMUNE | <20 | 751 | 26 | 429 | 3,00 | 57,53 |
| HPP 39 | 26 | F | ATROIZA | <20 | 953 | 25 | 497 | 4,76 | 38,9 |
| HPP 46 | 24 | F | TRIBUSS | <20 | 708 | 25 | 307 | 3,85 | 87,54 |
| HPP 50 | 27 | F | TRIBUS | <20 | 855 | 26 | 524 | 4,36 | 48,47 |
| HPP 66 | 22 | M | ATROIZA | <20 | 1103 | 14 | 624 | 3,72 | 10,12 |
| HPP 70 | 20 | F | ATROIZA | <20 | 729 | 19 | 385 | 4,04 | 68,87 |
| HPP 71 | 21 | M | TRIBUS | <20 | 734 | 20 | 473 | 4,28 | 14,04 |
| HPP 73 | 31 | M | ATROIZA | <20 | 711 | 42 | 565 | 3,36 | 42,04 |
| HPP 75 | 25 | F | ATROIZA | <20 | 927 | 25 | 573 | 3,65 | 47,08 |
| HPP 82 | 24 | M | ODIMUNE | <20 | 834 | 36 | 317 | 4,20 | 55,56 |
| HPP 84 | 25 | M | ODIMUNE | <20 | 643 | 19 | 371 | 2,72 | 45,42 |
| HPP 86 | 21 | F | ATROIZA | <20 | 1352 | 22 | 384 | 5,20 | 98,08 |
| HPP 91 | 26 | F | ATROIZA | <20 | 843 | 25 | 482 | 2,46 | 31,14 |
| HPP 104 | 24 | M | ODIMUNE | <20 | 252 | 54 | 199 | 3,71 | 32,32 |
| HPP 105 | 21 | F | ODIMUNE | <20 | 775 | 20 | 523 | 4,65 | 56,92 |
| HPP 109 | 25 | F | ATROIZA | <20 | 618 | 27 | 618 | 2,23 | 72,71 |
| HPP 118 | 32 | M | ATROIZA | <20 | 339 | 20 | 165 | 5,30 | 48,47 |
| **Elite controllers** |  |  |  |  |  |  |  |  |  |
| HPP 106 | 24 | F | ATROIZA | <20 | 782 | 33 | 612 | NA | 17,93 |
| HPP 128 | 21 | M | ATROIZA | <20 | 579 | 14 | 479 | NA | 68,4 |

## Discussion

Emerging work has demonstrated that Tat/Rev msRNA expression, which correlates with plasma HIV-1 RNA upon latency reversal, may be a relevant and promising indicator of the inducible, replication-competent viral reservoir; strong correlation was observed between supernatant HIV-1 RNA and msRNA but not usRNA upon latency reversal ex vivo, suggesting msRNA to be a more relevant biomarker of virion production^45^. Existing approaches to quantify the levels of Tat/Rev msRNA utilize real time RT-qPCR or RT-ddPCR on bulk total cellular RNA^39, 41^. These methods are highly sensitive and specific for the targeted viral RNA transcript. Certain RNA induction assays directly probe for Tat/Rev msRNA using whole cells, without RNA extraction, which may substantially simplify the workflow in that regard. However, semi-nested RT-qPCR aims to amplify the low copies of Tat/Rev msRNA transcripts to detectible and quantifiable levels^38, 48^. Furthermore, it should be noted that the direct whole cell input without DNase digestion followed by semi-nested RT-qPCR method of quantification as we and others have previously utilized in different studies^38, 63–66^ poses several limitations, with potentially direct consequences on the accuracy of the assay endpoint. An important limitation is the increased likelihood of detecting Tat/Rev within genomic HIV-DNA, which would introduce assay false positives, leading to overestimated measurements of the inducible viral reservoir. Moreover, semi-nested RT-qPCR approaches, performed in limiting dilution, are low throughput and considerably increase cross-contamination risks, the latter leading to false measurements. To address these two critical caveats of Tat/Rev msRNA RT-qPCR method of detection, we leveraged the superior qualities of RT-LAMP to develop SQuHIVLA, a rapid, sensitive, highly accurate and specific assay for Tat/Rev msRNA detection and quantification. SQuHIVLA detects and quantifies HIV-1-infected cells that spontaneously or inducibly express Tat/Rev msRNA and provides a first demonstration of LAMP technology for HIV-1 viral reservoir assessment and quantification.

The highly sensitive detection of Tat/Rev msRNA within half an hour of target amplification is revolutionary and superior to existing semi-nested RT-qPCR methods. We capitalized on this rapid amplification, high sensitivity and specificity of RT-LAMP, performed in a single reaction, to enhance the method of detecting and quantifying cells expressing Tat/Rev msRNA upon activation (SQuHIVLa). We have demonstrated, as a proof of concept, RT-LAMP-based inducible viral reservoir assessment in CD4+ T cells from a diverse group of people living with HIV-1 subtype B and C on suppressive antiretroviral therapy. Although limited in number, our data indicate good sensitivity, specificity and low inter-assay variability. With a reservoir size as low as 10 cells per million of CD4+ T cells, we were able to quantify the inducible reservoir of PLWH, proving the high sensitivity of our test. Further assessment of the assay using cells from individuals with low or ultra-low viral reservoir size is warranted.

In developing LAMP primers for the exclusive detection of Tat/Rev msRNA, it was essential to design primers that were exon-spanning. The Tat/Rev intron, approximately 2 kb, if incorporated into the forward or backward loop, during amplification, would decrease the stability of the formed loops, thereby inhibiting further target amplification and the fluorescent self-quenching loop primer^67^, which is used as a reporter for specific target amplification, would be non-functional. We have developed a comprehensive set of guidelines to follow when designing Tat/Rev exon-spanning LAMP primers, which must fulfil crucial target-specific criteria in addition to the standard LAMP primer requirements. Designing LAMP PCR primers and a probe to bind within the Tat/Rev genomic region is extremely challenging due to high HIV-1 genetic diversity and the need to identify eight relatively conserved primer binding regions for the LAMP reaction. A universal LAMP primer/probe set for the broad detection of Tat/Rev msRNA expressed from multiple HIV-1 subtypes is quite likely not feasible. It may be possible to multiplex LAMP primer/probe sets for different HIV-1 subtypes into one LAMP reaction. However, this will be challenging to achieve^68^ without complex molecular design^69, 70^ and will require an extensive number of optimizations.

Interestingly, in our cohort of PLWHC, treated during chronic infection stage, the mean frequency of cells expressing Tat/Rev msRNA after *in vitro* treatment with PMA/Ionomycin was significantly higher in women compared to men. Few studies have investigated sex differences in HIV-1 reservoir size, mostly determined by methods to quantify HIV-1 DNA and integrated HIV DNA, which have been reported to be comparable in men and women^71–73^. Studies that have assessed the viral transcription and replication by quantitating levels of cell-associated unspliced HIV-1 RNA and viral outgrowth have revealed a smaller inducible viral reservoir in women^74^, which contradicts our findings. This discrepancy however could be explained by differences in the endpoints, unspliced versus multiply spliced viral RNA transcripts measured by the different assays, which reflect different capacities of viral inducibility. Furthermore, reservoir quantification of viral outgrowth is often an underestimation due to suboptimal induction of majority of proviruses^75^. It is likely that the frequency of inducible, replication-competent proviruses is truly small in women, or poorly inducible. Indeed, HIV-1 inducibility in women, is potentially influenced by estrogen, which has been demonstrated to repress HIV-1 transcription in latency models^76^, in vitro infection systems^77^, and primary cells^76^, implicating a direct role for sex hormones mediating sex differences in reservoir maintenance and dynamics. Further research with a larger sample size and mechanistic studies are needed to better understand any potential sex-specific differences in the inducible reservoir size and the implications thereof.

Amid the rise of explorative strategies to clear the inducible viral reservoir in PLWH, reliable, sensitive and scalable assays are essential to determine the efficacy of putative intervention strategies. Moreover, it will be important for future studies to assess large cohorts in Sub-Saharan Africa^78^, which bears a disproportionate burden of HIV-1^2^. This also points to the need for the applicability of any future viral reservoir quantification assays in resource-constrained settings^79^.RT-LAMP is expected to contribute to this goal as it is highly sensitive, capable of detecting low copy numbers of RNA targets, and can provide quantitative results through real-time monitoring. Furthermore, it is less expensive, more user-friendly, and compatible with a variety of sample types. These features make RT-LAMP a promising tool for quantitative analysis in various fields, including HIV-1 molecular diagnostics and viral load monitoring especially in limited resource settings. Here, we have demonstrated that the exclusive detection of Tat/Rev msRNA, a proximal surrogate marker for inducible, replication-competent HIV-1, is possible using SQuHIVLa. Further research and collaboration with biomedical industries is needed to develop certified high-throughput assays for HIV-1 and other pathogens based on the principles of SQuHIVLa.

## Acknowledgments

T.M. received funding from the European Research Council (ERC) under the European Union’s Seventh Framework Programme (FP/2007–2013)/ERC STG 337116 Trxn-PURGE; Dutch Aidsfonds [2014021]; Health Holland [grants LSHM19100-SGF and EMCLSH19023]; ZonMW [grant 40-44600-98-333]; Erasmus MC mRACE research grant. R.J.P received funding from Dutch Aidsfonds [2016014]; S.R. received funding from Dutch Aidsfonds [P-53302]. P.M. received funding from South African Medical Research Council.

## Author contributions

Conceptualization, T.H., C.L. and T.M.; Methodology, T.H., C.L. and T.M.; Investigation, T.H., C.L., S.D.S., M.K., A.N. and T.W.K.; Funding Acquisition, T.M., S.R., R.J.P, P.M. and T.N.; Project Administration, T.M., S.R., R.J., P.M. and T.N.; Supervision, T.M. and T.N. Writing – Original Draft, T.H., C.L., S.D.S, M.K. and T.M.; Writing – Review & Editing, T.H., C.L., S.D.S, M.K., S.R., A.N., T.W.K., R.J.P., P.M., T.N., and T.M.

## Methods

### Cohort characteristics

Peripheral blood samples from 25 people living with HIV-1 (PLWH), 3 subtype B and 19 subtype C were used in this study. All 22 PLWH included in this study were older than 18 years of age and on fully suppressive combination antiretroviral therapy (cART), initiated during chronic stages of HIV-1 infection, with plasma HIV-1 RNA below detectable levels (< 20 copies/mL) for at least twelve months. Additional clinical information is shown in Table 1.

### Ethical statement

Written informed consent was obtained from all study participants. The Medical Ethical Council of the Erasmus Medical Center (MEC-2012–583) and University of KwaZulu-Natal (REF: E036/06) approved the use of clinical material for research purposes.

### Sample collection

Peripheral blood mononuclear cells (PBMCs) were isolated from whole blood or leukapheresis samples using Ficoll density gradient centrifugation and cryopreserved in liquid nitrogen until further use. CD4+ T cells were isolated from thawed PBMCs by negative magnetic selection using the EasySep Human CD4 T Cell Enrichment Kit (STEMCELL Technologies) according to the manufacturer’s protocol.

### Designing of Tat/Rev RNA-specific LAMP primer/probe sets

Briefly, ten complete HIV-1 genome sequences of a particular subtype, submitted from various geographical locations and from different years, were obtained from the Los Alamos HIV sequence databases (https://www.hiv.lanl.gov/) (GenBank Accession numbers of the sequences used to design subtype B and C Tat/Rev msRNA LAMP primers are listed in supplemental table S1). The sequences were aligned using the Molecular Evolutionary Genetics Analysis software version 11 (MEGA11)^80^ and manually modified to generate a new alignment of *in silico*-spliced Tat/Rev DNA sequences (approximately 490 bp). The consensus Tat/Rev DNA sequence of this second alignment was then converted to a FASTA file and used to generate preliminary sets of LAMP primers using the online program PrimerExplorer V5 (http://primerexplorer.jp/e/). Selected LAMP primers, based on criteria that we have identified to be crucial to ensure precise detection of Tat/Rev msRNA transcripts, were aligned to the complete HIV-1 genome alignment to assess primer-target sequence complementarity. To broaden the detection of multiple strains within a particular HIV-1 subtype, the primer binding regions with mutations present at sites extremely important for amplification were avoided by shifting these regions several nucleotides up or downstream. Importantly, the melting temperature of the new primers and the distance between the primer binding regions were re-calculated to ensure that the modifications still met the initial LAMP primer specifications. Additionally, loop primers were generated using the PrimerExplorer V5 software and one of these (e.g., Loop forward primer) was converted into a self-quenching probe with an internal FAM fluorophore bound to the closest thymidine from the 3’ end^67^. A more detailed protocol for designing these HIV-1 Tat/Rev msRNA LAMP primers and probe is provided below:

### Steps to design HIV-1 Tat/Rev msRNA LAMP primers

#### 1) Select reference HIV-1 genome sequences

To design LAMP primers for a specific subtype of HIV-1, download a minimum of 10 complete genome sequences of the desired subtype from the Los Alamos database using the sequence search interface webpage (https://www.hiv.lanl.gov/components/sequence/HIV/search/search.html). Select sequences submitted from different geographic locations (such as, Africa, Asia, Europe and USA) and in different years (e.g., sequences submitted before 2000, between 2000-2010 and 2010-present). This selection is necessary to account for sequence diversity and increase the probability that the designed primers will recognize and bind to Tat/Rev msRNA from a majority of individuals with the selected HIV-1 subtype. Download the selected reference HIV-1 genome sequences, save in FASTA file format.

#### 2) Generate a spliced Tat/Rev consensus sequence *in silico*

Align the downloaded reference HIV-1 genome sequences using a sequence alignment tool of choice. We utilized the built-in MUSCLE algorithm in MEGA11 software. Manually edit the aligned sequences by excluding the following sequence regions; upstream of the Tat gene (1 to 5830); Tat/rev intron (6046 to 8383) and downstream of the *rev* gene (8654 to 3’end). The nucleotide positions are relative to the coordinates of the 5’ LTR UR start of the reference HIV-1 genome (ID) provided in the Los Alamos HIV-1 sequence compendium (2021). The resulting alignment of the spliced Tat/Rev sub-genomic region should be approximately 490 nucleotides. Download the spliced Tat/Rev DNA consensus sequence, save in FASTA file format.

### Generate a preliminary LAMP primer set using the PrimerExplorer V5 software

Upload the Tat/Rev DNA consensus FASTA file (i.e., the target sequence file) into the online PrimerExplorer V5 software. Adjust the default settings prior to executing the primer design algorithm. Set the sequence parameter condition to “AT rich”; the GC content of the Tat/rev sequence varies from 35-45% depending on the HIV-1 subtype. Set the Tm for F1c/B1c to 60°C-64°C, and set the Tm for F3/B3 and F2/B2 to 55°C-60°C to ensure the generation of a sufficient number of potential LAMP primer sets. The software displays a maximum of 1000 primer sets using a combination of different primer binding sites.

Several prerequisites need to be considered when selecting the preliminary primer set from the potential LAMP primer sets generated by the software.

A) To ensure the specificity of the primer set for Tat/Rev msRNA and not the intron-containing Tat/Rev DNA, only primer sets containing the splice site of msRNA (approximately, the 215^th^ nucleotide) within the F2 to F1c or B2 to B1c primer binding regions should be considered. This ensures that unstable loop formation, while the primers are bound to intron-containing Tat/Rev DNA, would result in the inhibition of isothermal amplification. Therefore, select potential primer sets where the F2 binding region is within 25 nucleotides downstream of the splice site and 90 nucleotides upstream of the splice site (approximately, nt pos. 125 to 240) or where the B2 binding region is within 25 nucleotides upstream of the splice site and 90 nucleotides downstream of the splice site (approximately, nt pos. 190 to 305).
B) Since loop primers, which accelerate the LAMP reaction, are also used for amplification, only primer sets that have a distance of >22 nucleotides between F2 and F1c/B2 and B1c should be considered for the preliminary primer set.
C) If impossible to find one primer set that fulfills both requirements for exclusive binding of msRNA, and suitable for designing the loop primers, then primer binding sites belonging to different primer sets could be mixed to form a custom set as long as they have a similar Tm for F3/B3, F2/B2, and F1c/B1c and fulfill the specific distance requirement between LAMP primer binding regions.

After selecting the preliminary primer set, download the sequences and primer information files from the software to use for designing the Loop primers (LF/LB).

#### 3) Design loop primers

Upload the LAMP primer information files into the PrimerExplorer V5 software. Since one of the loop primers needs to be converted into a self-quenching and internal FAM fluorophore-containing probe, specific criteria should be considered when selecting the loop primers:

A) presence of a cytosine (C) or guanine (G) residue at the terminal 3′ end
B) a thymine (T) residue at the second or third position from this 3′ end
C) one or more G nucleotides flanking the T residue (optional).

In case no suitable loop primers can be selected from the software-generated list, the closest possible sequence is chosen, and manual modifications are performed to ensure its applicability as a probe (detailed below).

#### 4) Adapt LAMP primer and probe sequences

The Tat/Rev region of HIV-1 is highly diverse, making it difficult to find completely conserved primer binding regions for LAMP primers. To address this, align the primer binding regions of the preliminary primer set selected in **step 3** to the reference HIV-1 sequences downloaded from the Los Alamos database in **step 1**. Identify mutation hotspots located within primer binding regions that would result in primer-template mismatches. Ensure that the primers contain conserved nucleotides at specific positions that are essential for amplification to proceed (3’ end of F2/B2 and F3/B3, and 5’ end of F1c/B1c. Mutation hotspots are permitted at less essential regions (5’ end of F2/B2 and F3/B3, 5’ end of the F1c/B1c, and the internal region) where a mismatch will have less effect on amplification. To ensure these criteria, shift the preliminary primers upstream or downstream of the current primer binding site. Recalculate the Tm for each modified primer to ensure that the new primers have similar Tm to the original primers, and that the distance between the modified primer regions must also meet LAMP-specific requirements. The distances between the primer binding regions are calculated from the alignment and the Tm of the modified primers is calculated using Kun’s Oligonucleotide Tm calculator (https://arep.med.harvard.edu/kzhang/cgi-bin/myOligoTm.cgi). The calculated Tm is affected by experimental conditions such as the salt concentration and oligo concentration, so it is preferred that Tm be calculated under fixed experimental conditions (oligo concentration at 0.1 µM, sodium ion concentration at 50 mM, magnesium ion concentration at 4 mM).

Similarly, align the selected loop primers to the reference HIV-1 complete genome sequences and modify following the same procedures (shifting the loop primers upstream or downstream of the current primer binding site). Ensure that one of the loop primers fulfills the requirements to be converted into a self-quenching probe and that the internally labeled T and 3’ end G/C of the probe region are conserved among the HIV-1 sequences used for the alignment.

The sequences of the HIV-1 genotype B and C LAMP primers and probes designed and used to validate specific detection of Tat/Rev msRNA by RT-LAMP are provided in Supplemental Table S2.

### RT-LAMP assay validation using *in vitro* Tat/Rev RNA transcripts

A gBlock (IDT) comprising a spliced Tat/Rev sequence (subtype B or C), downstream of a T7 promoter sequence (generated by Integrated DNA Technologies) was used as a template for in vitro transcription using Hi-T7 RNA Polymerase and Ribonucleotide Solution Mix (both from New England Biolabs) in accordance with the manufacturer’s instructions. The sequences of the subtype B and subtype C-specific Tat/Rev gBlocks are listed in Supplemental table S3. To eliminate gBlock DNA, in vitro-transcribed Tat/Rev RNA samples were treated with RNase-free DNase I (New England Biolabs) and purified using the Monarch® RNA Cleanup Kit (New England Biolabs). Purified Tat/Rev RNA samples were serially diluted to achieve concentrations of 1000, 500, 250, 125, 50, 20, 10, 5, 1, 0.1 copies of RNA/ 5 μL and used in validation experiments to test the sensitivity of the RT-LAMP reactions. A single RT-LAMP reaction consisted of enzymes (RTx (NEB), Bst 2.0 (NEB), RNasin (Promega), Tat/Rev RNA primer/probe set, and Tat/Rev RNA template to a final reaction volume of 20 μL. Reactions without the RTx enzyme were performed to assess the specificity of the RT-LAMP assay in exclusively amplifying RNA template. The amplification was carried out in a CFX96 Touch Real-Time PCR Detection System thermocycler (BioRad) following a thermal program of continuous 65°C with fluorescence read every 30 s for 180 cycles, which corresponds to 90 min of amplification time. Complete details of the RT-LAMP reagents and conditions for reaction are listed in Supplemental table S4. Positive reactions were identified by the presence of fluorescence curves exceeding the cutoff line.

### RT-LAMP-based detection of Tat/Rev msRNA in J-Lat 11.1 cells

Latent HIV-1-infected Jurkat cells (clone J-Lat 11.1), which harbor a full integrated HIV-1 subtype B genome with a mutated *env* gene and a GFP reporter gene in place of *nef*, which requires multiple splicing (Figure S1C) ^81^, served as a surrogate for inducible reservoir cells. Single J-Lat 11.1 cells in the background of uninfected donor CD4+ T cells were used for the validation of RT-LAMP-based detection of Tat/Rev msRNA. J-Lat 11.1 cells were cultured in complete RPMI-1640 media, supplemented with 7% FBS and 100 μg/ml penicillin-streptomycin, at 37°C in a humidified 5% CO_2_ incubator. Primary CD4+ T cells isolated from uninfected donor PBMCs were cultured in RPMI-1640 media supplemented with 10% FBS and 100 μg/ml penicillin-streptomycin at 37°C in a humidified 5% CO_2_ incubator. J-Lat 11.1 cells were stimulated with 10 μM of phorbol 12 myristate 13-acetate (PMA) (Sigma) for 12 hrs and primary CD4+ T cells were stimulated with 100 ng/mL of PMA (Sigma) and 1 µg/mL of ionomycin (Sigma) for 12 hrs. A single GFP+ J-Lat 11.1 cell (marking a Tat/Rev msRNA+ cell) was sorted directly into each well of a 96 well white PCR plate (BioRad) containing the RT-LAMP master mix, described in Supplemental table S4, without cells or with an increasing background of activated uninfected donor CD4+ T cells (5x10^3^, 1x10^4^, 2x10^4^, 4x10^4^, 8x10^4^ cells per reaction). PCR plates were then sealed and RT-LAMP was carried out using a CFX96 Touch Real-Time PCR Detection System thermocycler (BioRad) following a thermal program of continuous 65°C with fluorescence read every 30 s for 180 cycles, which corresponds to 90 min of amplification time. Positive reactions were identified by the presence of fluorescence curves exceeding the cutoff line.

### Validating specificity for Tat/Rev HIV-1 msRNA by RT-LAMP

After 12 hr stimulation of J-Lat 11.1 with PMA (see above) either a single GFP+ or a single GFP-J-Lat 11.1 cell was sorted directly into each well of a 96 well PCR plate containing complete RT-LAMP master mix or the master mix without the reverse transcriptase (RT). PCR plates were then sealed and amplification was carried out with the CFX96 Touch Real-Time PCR Detection System thermocycler (BioRad) following a thermal program of continuous 65°C with fluorescence read every 30 s for 180 cycles, which corresponds to 90 min of amplification time. In order to compare the specificity of RT-LAMP that utilizes exon-spanning msRNA specific LAMP primers/probe with semi-nested RT-qPCR that utilizes non exon-spanning primers/probe reported in previous study^38^, a single GFP+ or GFP-J-Lat 11.1 cell was sorted directly into each well of a 96-well PCR plate containing the pre-amplification mastermix consisting of 1 μL One-step RT-PCR enzyme (QIAGEN), 5 μL 5× One-step RT-PCR buffer (QIAGEN), 10 μL Triton-x100 (0.3%) (Sigma), 0.25 μL RNAsin (40 U/μL) (Promega), 1 μL dNTPs (10 mM each), 0.5 μL of forward tat 1.4 and reverse rev primers^38^ (both at 20 µM), and nuclease-free water to a final volume of 25 µL. The PCR plates were sealed, and T100 thermocycler (BioRad) was used to carry out the following thermal program; reverse transcription at 50 °C for 30 min, denaturation at 95 °C for 15 min, followed by 25 cycles of 95 °C for 30 s, 55 °C for 1 min and 72 °C for 2 min, and a final extension at 72 °C for 5 min. Immediately following pre-amplification, 2 µL of the 1st PCR product was directly used as input for the Tat/Rev semi-nested real-time PCR reaction, which included 5 µL 4× Taqman Fast Advanced Master Mix (Thermo Fisher Scientific), 0.4 µL of each primer (tat 2.0 and rev, both at 20 µM), 0.4 µL probe (5 µM)^38^ and nuclease-free water to a final reaction volume of 20 µL. The following thermal settings were used for the final amplification on a BioRad CFX96 Touch Real-Time PCR Detection System thermocycler: 5 min at 50 °C (UNG step), 95 °C for 20 s, followed by 45 cycles of 95 °C for 3 s and 60 °C for 30 s.

Genomic DNA was isolated from PMA-stimulated J-Lat 11.1 cells using the phenol–chloroform-isoamyl alcohol isolation method and ethanol precipitation in the presence of glycogen as a carrier. Isolated DNA was treated with DNase-free Monarch® RNase A (New England Biolabs) to get rid of any RNA contamination. Total RNA was also isolated from PMA-stimulated J-Lat 11.1 cells using Trizol reagent (Sigma) according to the manufacturer’s instructions. Isolated RNA was treated with RNase-free DNase I to get rid of any DNA contamination. 100 ng of DNase-treated RNA, 100 ng of RNase-treated DNA and 100 copies HIV-1 plasmid pNL4-3.Luc.R-E-were used as template to perform semi-nested RT-qPCR (following above mentioned procedure) and RT-LAMP on a CFX96 Touch Real-Time PCR Detection System thermocycler (BioRad) following a thermal program of continuous 65°C with fluorescence read every 30 s for 180 cycles, which corresponds to 90 min of amplification time. Positive reactions were identified by the presence of fluorescence curves exceeding the cutoff line.

### SQuHIVLa validation for viral reservoir quantification

To perform SQuHIVLa in custom samples representative of clinical samples with an inducible HIV-1 subtype B reservoir of 0.1, 1, 10, 20 cells/million CD4+ T cells, J-Lat 11.1 cells and CD4+T cells derived from uninfected donor PBMCs were cultured and stimulated following the same procedure mentioned in the previous section. After stimulation, CD4+ T cells were washed and counted using an automated cell counter (Countess II, ThermoFisher). A range of 1, 10, 100 or 200 GFP+ J-Lat 11.1 cells were sorted into separate tubes containing activated, uninfected donor CD4+ T cells in RPMI-1640 3% FBS (1 million CD4+ cells/mL). The cells in each tube were then washed, resuspended in phosphate-buffered saline (PBS) and then serially diluted to 4 × 10^6^ cells/ml, 2 × 10^6^ cells/ml, 1 × 10^6^ cells/ml and 5 × 10^5^ cells/ml in PBS. From each dilution, 5 µL of the cell suspension was distributed to 22–24 wells of a 96-well plate containing 15 µL RT-LAMP master mix (Supplemental table S4) corresponding to 20000, 5000, 1250 and 313 cells per well. After the RT-LAMP Reaction, the positive wells at each dilution were scored, and the maximum likelihood method was used to determine the frequency of cells expressing Tat/Rev msRNA using the IUPMStats v1.0 online software^82^.

Additional SQuHIVLa validation experiments were performed using peripheral blood samples from people living with HIV (PWH). Primary CD4+ T cells isolated from thawed PBMCS were resuspended in culture RPMI-1640 media supplemented with 10% FBS and 100 μg/ml penicillin-streptomycin and rested for five hrs at 37°C in a humidified, 5% CO_2_ incubator. CD4+T cells were then stimulated with 100 ng/mL of PMA and 1 µg/mL of ionomycin (both from Sigma) for 12 hrs. Activated CD4+ T cells were washed in RPMI 1640 media supplemented with 3% FBS, counted at least twice using Countess II (ThermoFisher) and serially diluted to 4 × 10^6^ cells/ml, 1 × 10^6^ cells/ml, 2.5 × 10^5^ cells/ml and 6.3 × 10^4^ cells/ml in PBS. From each dilution, 5 µL of the cell suspension was distributed to 22–24 wells of a 96-well plate containing 15 µL RT-LAMP master mix (Supplemental table S4). PCR plates were then sealed and RT-LAMP was carried out using a CFX96 Touch Real-Time PCR Detection System thermocycler (BioRad) following a thermal program; incubation at 45°C for 60 mins (determined based on the results depicted in Figure 3E) followed by continuous 65°C with fluorescence read every 30 s for 180 cycles, which corresponds to 90 min of amplification time. After the RT-LAMP reaction, the positive wells at each dilution were scored, and used to determine the frequency of cells expressing Tat/Rev msRNA, using the IUPMStats v1.0 online software^82^, which uses maximum likelihood statistics.

When a limited number of PBMCs were available, activated CD4+ T cells were serially diluted to 2 × 10^6^ cells/ml, 1 × 10^6^ cells/ml, 2.5 × 10^5^ cells/ml and 6.3 × 10^4^ cells/ml in PBS and, 5 µL of the cell suspension was distributed to 22–24 wells of a 96-well plate corresponding to 10000, 5000, 1250 and 313 cells per well.

### Statistical analysis

All graphs were generated and statistical analyses were performed using GraphPad Prism version 8.0.2 for Windows (GraphPad Software, San Diego, California USA, www.graphpad.com). In order to calculate the lowest amount of Tat/Rev msRNA in a sample that can be consistently detected with 95% probability (LOD 95%), probit analysis was used. The Probit was calculated by the Excel function [5+NORMSINV(P)], where P was the hit-rate (proportion of positive reactions) and 6.64 probit value (1.64 for 95% limit, +5 for probit scale) was used to extrapolate LOD 95% RNA copies. The effect of background cells on RT-LAMP amplification time was determined using Ordinary One-way ANOVA followed by Dunnett’s multiple comparisons test. Coefficient of variation (CV) was calculated to determine the reproducibility of SQuHIVLa assay. Normality of the calculated inducible HIV-1 reservoir size (Figure 4D) using SQuHIVLa was tested with the Shapiro-Wilk test. All the data passed the normality test which is why correlations were performed using the Pearson test. Accuracy percentage (AP) in Figure 4D was calculated using ABS function in Excel. Inducible HIV-1 subtype C reservoir size quantified from infected individuals with different biological sex were compared utilizing two-tailed unpaired t-test.

## Supplemental figures

**Figure S1. Tools used for evaluating RT-LAMP sensitivity.** A) Percentage of positive LAMP reactions and B) amplification time required for different amount of RNA copies are plotted when the amplification was carried out without reverse transcriptase enzyme. Data are presented as mean ± SD of three independent reactions, each of which had four technical replicates. C) Schematic overview of the integrated HIV-1 genome in JLat 11.1 cells. The green rectangle depicts the substitution of the Nef gene with GFP in the integrated HIV-1 genome, and the red rectangle represents the mutation in the HIV-1 Env gene that results in a faulty viral envelope protein. D) Sorting of GFP+ positive JLat 11.1 cells upon PMA stimulation. GFP-positive cells are represented by the green cell population, and GFP-negative cells by the purple cell population. Related to figure 3.

**Figure S2: Comparison of specificity in detecting of msRNA.** A) Percentages of positive reactions when a single GFP+ or GFP-JLat 11.1 cell/reaction was used as input for RT-LAMP and Semi-nested PCR amplification. Data are presented as mean ± SD of three independent experiments, each of which included 15 technical replicates. B) Percentages of positive reactions when DNase treated RNA and RNase treated DNA sample isolated from PMA stimulated JLat 11.1 cells along with pNL4.3 E-R-plasmid were used as input for RT-LAMP and Semi-nested PCR amplification. C) Schematic overview of the HIV-1 Tat/Rev specific TILDA primers and probe binding sites. Related to figure 4.

**Figure S3: RT-LAMP for the precise detection of msRNA.** A) Amplification time required for RT-LAMP when a single GFP+ JLat 11.1 cell/reaction were used as input for RT-LAMP amplification with or without reverse transcriptase enzyme B) Amplification time required for RT-LAMP when a single GFP+ or GFP-JLat 11.1 cell/reaction were used as input for amplification. Data are presented as mean ± SD of three independent experiments, each of which had 15 technical replicates. Related to figure 4.

**Figure S4: Heat map of the primer mismatches.** Heat maps of the mismatches of subtype B Tat/Rev Primers binding to HIV-1 subtype B (A), C (B) and A (C). Y-axis represents different primers and x-axis represents nucleotide positions. The value, and corresponding color, in the heat map portrays the relative amount of mismatches that occur at each position. Related to figure 5.

**Figure S5: Correlation of inducible reservoir size of male and female participants quantified by SQuHIVLa with different clinical parameters.** The pearson correlation coefficient (r2) is determined between inducible reservoir size quantified using SQuHIVLa and CD4 nadir for 10 female participants (A) and 7 male participants (B). The pearson correlation coefficient (r2) is also determined between inducible reservoir size quantified using SQuHIVLa and CD4+ T cell count at the time of sampling for 10 female participants (C) and for 7 male participants (D). Female participants are depicted with blue open circle and male participants are depicted with red square. Statistical significance is determined by p < 0.05. Related to figure 5.

## Supplemental tables

**Supplemental Table S1:**
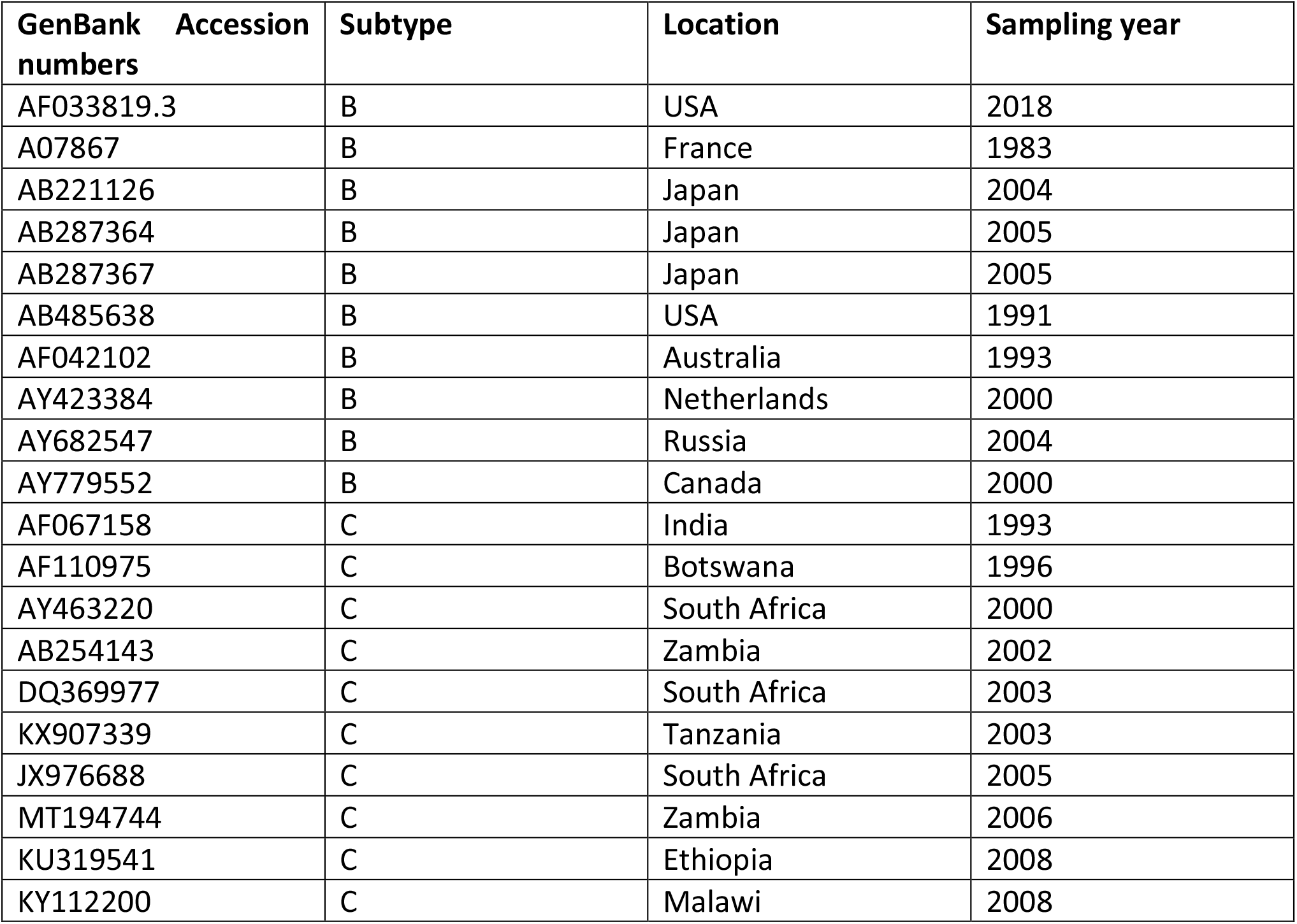
GenBank accession numbers of the sequences used to design Tat/Rev HIV-1 msRNA specific LAMP primers and probes. Related to figure 2 and 5.

**Supplemental Table S2:**
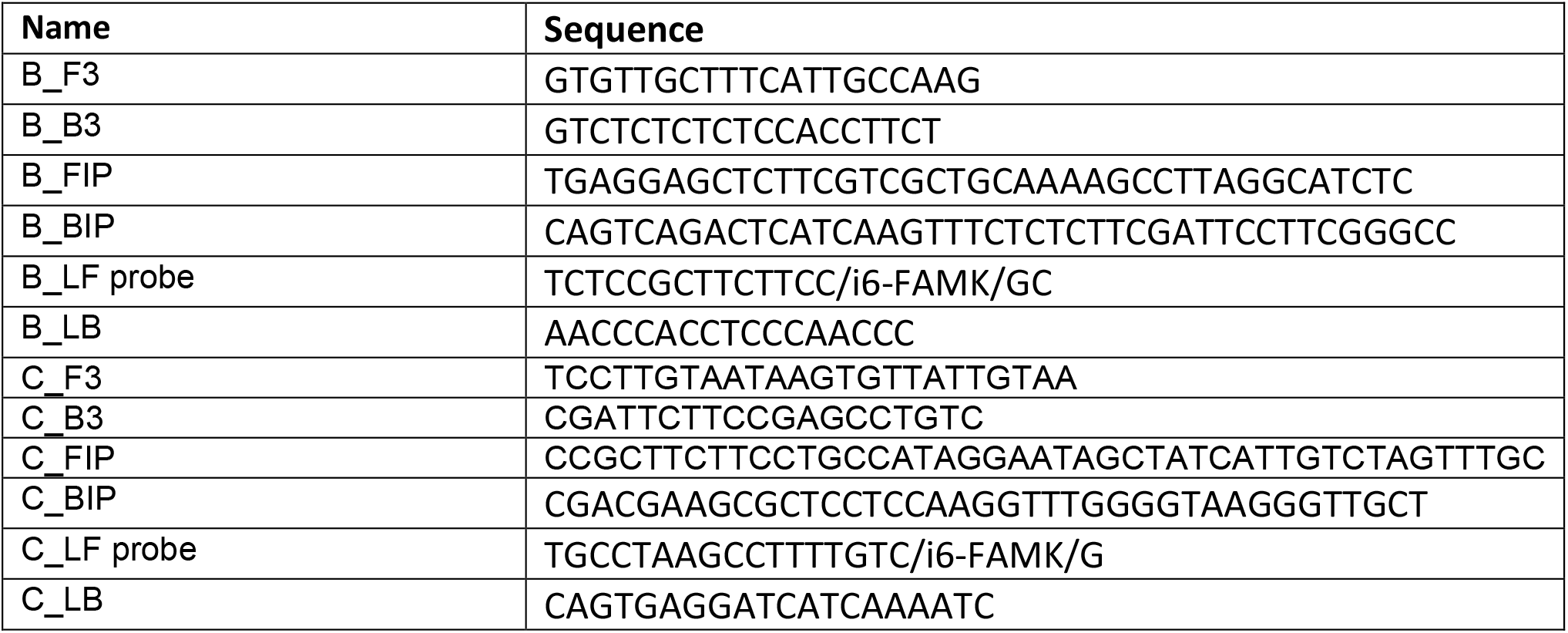
Tat/Rev HIV-1 msRNA specific LAMP primers and probes sequences. Related to figure 2, 3, 4 and 5.

**Supplemental Table S3:**
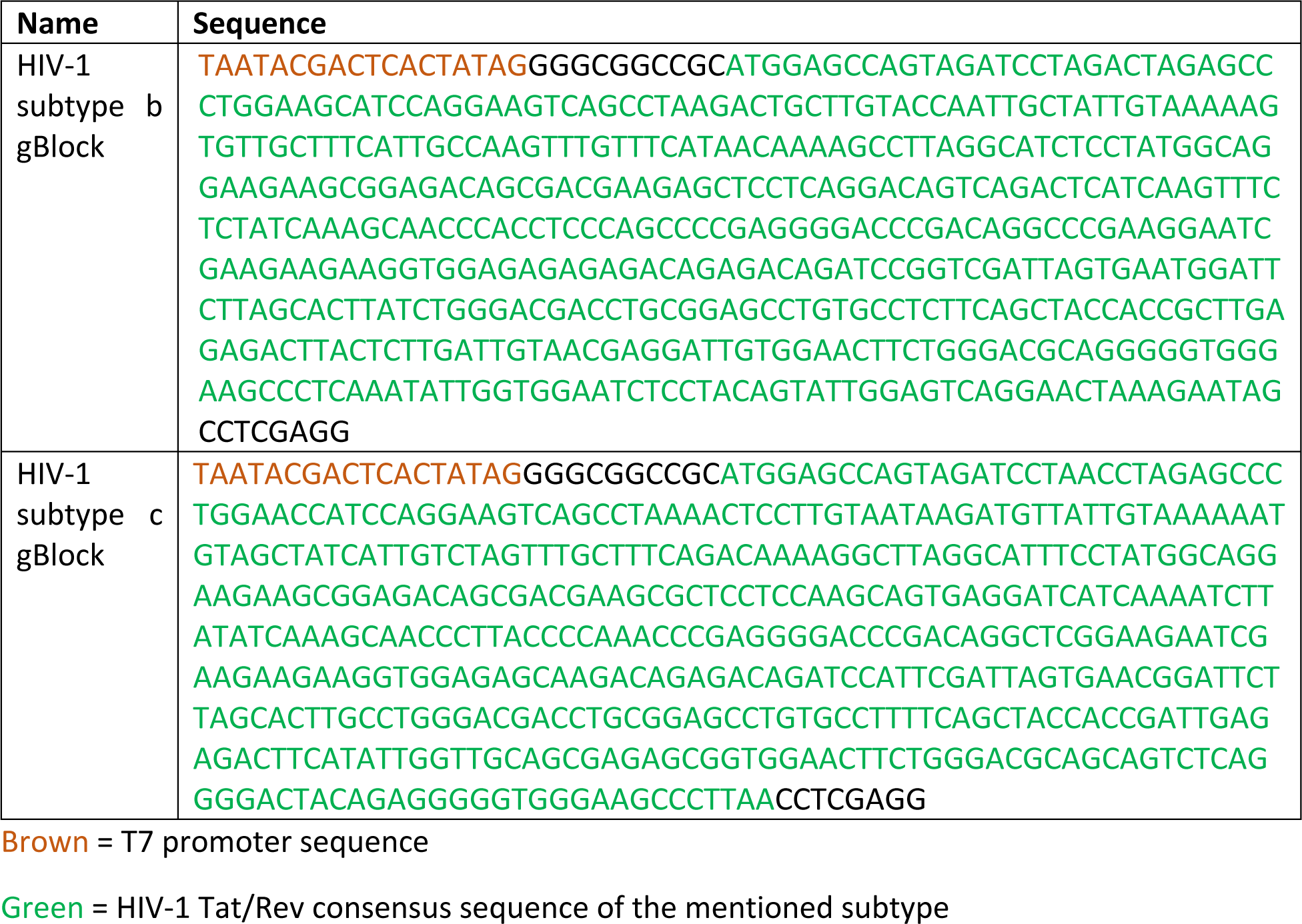
Tat/Rev HIV-1 gBlock sequences. Related to figure 3 and 5.

**Supplemental Table S4:**
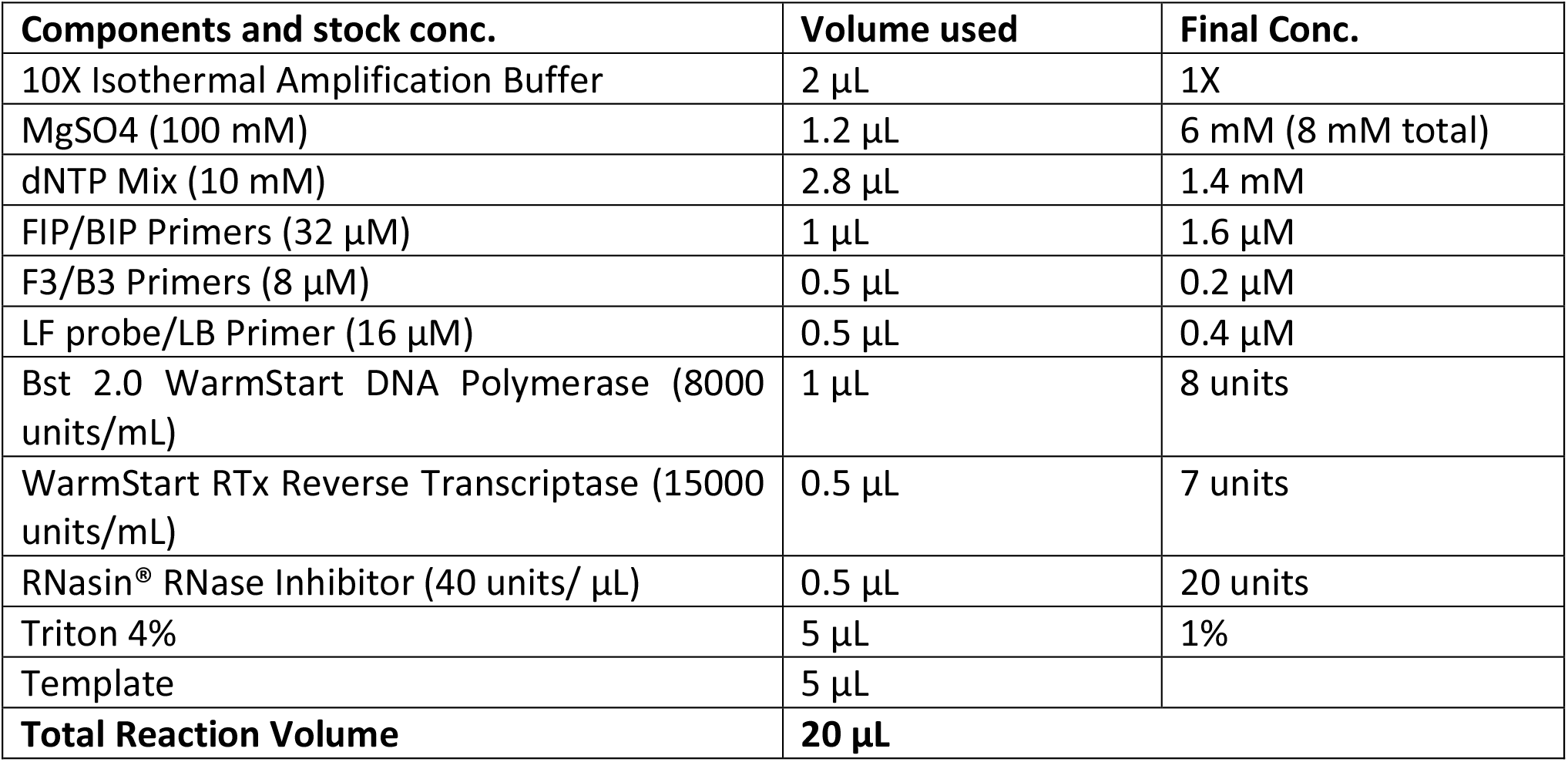
RT-LAMP mastermix composition. Related to figure 3, 4 and 5.

| **Components and stock conc.** | **Volume used** | **Final Conc.** |
| --- | --- | --- |
| 10X Isothermal Amplification Buffer | 2 μL | 1X |
| MgSO<sub>4</sub> (100 mM) | 1.2 μL | 6 mM (8 mM total) |
| dNTP Mix (10 mM) | 2.8 μL | 1.4 mM |
| FIP/BIP Primers (32 μM) | 1 μL | 1.6 μM |
| F3/B3 Primers (8 μM) | 0.5 μL | 0.2 μM |
| LF probe/LB Primer (16 μM) | 0.5 μL | 0.4 μM |
| Bst 2.0 WarmStart DNA Polymerase (8000 units/mL) | 1 μL | 8 units |
| WarmStart RTx Reverse Transcriptase (15000 units/mL) | 0.5 μL | 7 units |
| RNasin® RNase Inhibitor (40 units/ μL) | 0.5 μL | 20 units |
| Triton 4% | 5 μL | 1% |
| Template | 5 μL | |
| **Total Reaction Volume** | **20 μL** |  |

## References

1. Kharsany, A.B., and Karim, Q.A. (2016). HIV Infection and AIDS in Sub-Saharan Africa: Current Status, Challenges and Opportunities. Open AIDS J 10, 34–48.

2. UNAIDS (2023). UNAIDS Technical Support Mechanism.

3. Chun, T.W., Carruth, L., Finzi, D., Shen, X., DiGiuseppe, J.A., Taylor, H., Hermankova, M., Chadwick, K., Margolick, J., Quinn, T.C., et al. (1997). Quantification of latent tissue reservoirs and total body viral load in HIV-1 infection. Nature 387, 183–188.

4. Chun, T.W., Stuyver, L., Mizell, S.B., Ehler, L.A., Mican, J.A., Baseler, M., Lloyd, A.L., Nowak, M.A., and Fauci, A.S. (1997). Presence of an inducible HIV-1 latent reservoir during highly active antiretroviral therapy. Proc Natl Acad Sci U S A 94, 13193–13197.

5. Joos, B., Fischer, M., Kuster, H., Pillai, S.K., Wong, J.K., Böni, J., Hirschel, B., Weber, R., Trkola, A., Günthard, H.F., and Swiss, H.I.V.C.S. (2008). HIV rebounds from latently infected cells, rather than from continuing low-level replication. Proc Natl Acad Sci U S A 105, 16725–16730.

6. Landovitz, R.J., Scott, H., and Deeks, S.G. (2023). Prevention, treatment and cure of HIV infection. Nature Reviews Microbiology. 10.1038/s41579-023-00914-1.

7. Stoszko, M., Ne, E., Abner, E., and Mahmoudi, T. (2019). A broad drug arsenal to attack a strenuous latent HIV reservoir. Curr Opin Virol 38, 37–53.

8. Stone, M., Rosenbloom, D.I.S., Bacchetti, P., Deng, X., Dimapasoc, M., Keating, S., Bakkour, S., Richman, D.D., Mellors, J.W., Deeks, S.G., et al. (2021). Assessing the Suitability of Next-Generation Viral Outgrowth Assays to Measure Human Immunodeficiency Virus 1 Latent Reservoir Size. J Infect Dis 224, 1209–1218.

9. Hodel, F., Patxot, M., Snäkä, T., and Ciuffi, A. (2016). HIV-1 latent reservoir: size matters. Future Virol 11, 785–794.

10. Siliciano, J.D., and Siliciano, R.F. (2018). Assays to Measure Latency, Reservoirs, and Reactivation. Curr Top Microbiol Immunol 417, 23–41.

11. Wang, Z., Simonetti, F.R., Siliciano, R.F., and Laird, G.M. (2018). Measuring replication competent HIV-1: advances and challenges in defining the latent reservoir. Retrovirology 15, 21.

12. Cicilionytė, A., Berkhout, B., and Pasternak, A.O. (2021). Assessing proviral competence: current approaches to evaluate HIV-1 persistence. Curr Opin HIV AIDS 16, 223–231.

13. Eriksson, S., Graf, E.H., Dahl, V., Strain, M.C., Yukl, S.A., Lysenko, E.S., Bosch, R.J., Lai, J., Chioma, S., Emad, F., et al. (2013). Comparative analysis of measures of viral reservoirs in HIV-1 eradication studies. PLoS Pathog 9, e1003174.

14. Alidjinou, E.K., Bocket, L., and Hober, D. (2015). Quantification of viral DNA during HIV-1 infection: A review of relevant clinical uses and laboratory methods. Pathol Biol (Paris) 63, 53–59.

15. Bruner, K.M., Murray, A.J., Pollack, R.A., Soliman, M.G., Laskey, S.B., Capoferri, A.A., Lai, J., Strain, M.C., Lada, S.M., Hoh, R., et al. (2016). Defective proviruses rapidly accumulate during acute HIV-1 infection. Nature Medicine 22, 1043–1049. 10.1038/nm.4156.

16. Ho, Y.C., Shan, L., Hosmane, N.N., Wang, J., Laskey, S.B., Rosenbloom, D.I., Lai, J., Blankson, J.N., Siliciano, J.D., and Siliciano, R.F. (2013). Replication-competent noninduced proviruses in the latent reservoir increase barrier to HIV-1 cure. Cell 155, 540–551.

17. Bruner, K.M., Wang, Z., Simonetti, F.R., Bender, A.M., Kwon, K.J., Sengupta, S., Fray, E.J., Beg, S.A., Antar, A.A.R., Jenike, K.M., et al. (2019). A quantitative approach for measuring the reservoir of latent HIV-1 proviruses. Nature 566, 120–125. 10.1038/s41586-019-0898-810.1038/s41586-019-0898-8 [pii].

18. Falcinelli, S.D., Ceriani, C., Margolis, D.M., and Archin, N.M. (2019). New Frontiers in Measuring and Characterizing the HIV Reservoir. Frontiers in Microbiology 10. 10.3389/fmicb.2019.02878.

19. Levy, C.N., Hughes, S.M., Roychoudhury, P., Reeves, D.B., Amstuz, C., Zhu, H., Huang, M.L., Wei, Y., Bull, M.E., Cassidy, N.A.J., et al. (2021). A highly multiplexed droplet digital PCR assay to measure the intact HIV-1 proviral reservoir. Cell Rep Med 2, 100243.

20. Cassidy, N.A.J., Fish, C.S., Levy, C.N., Roychoudhury, P., Reeves, D.B., Hughes, S.M., Schiffer, J.T., Benki-Nugent, S., John-Stewart, G., Wamalwa, D., et al. (2022). HIV reservoir quantification using cross-subtype multiplex ddPCR. iScience 25, 103615.

21. Gaebler, C., Lorenzi, J.C.C., Oliveira, T.Y., Nogueira, L., Ramos, V., Lu, C.L., Pai, J.A., Mendoza, P., Jankovic, M., Caskey, M., and Nussenzweig, M.C. (2019). Combination of quadruplex qPCR and next-generation sequencing for qualitative and quantitative analysis of the HIV-1 latent reservoir. J Exp Med 216, 2253–2264.

22. Gaebler, C., Falcinelli, S.D., Stoffel, E., Read, J., Murtagh, R., Oliveira, T.Y., Ramos, V., Lorenzi, J.C.C., Kirchherr, J., James, K.S., et al. (2021). Sequence Evaluation and Comparative Analysis of Novel Assays for Intact Proviral HIV-1 DNA. J Virol 95.

23. Kinloch, N.N., Ren, Y., Conce Alberto, W.D., Dong, W., Khadka, P., Huang, S.H., Mota, T.M., Wilson, A., Shahid, A., Kirkby, D., et al. (2021). HIV-1 diversity considerations in the application of the Intact Proviral DNA Assay (IPDA). Nature Communications 12, 165. 10.1038/s41467-020-20442-3.

24. White, J.A., Kufera, J.T., Bachmann, N., Dai, W., Simonetti, F.R., Armstrong, C., Lai, J., Beg, S., Siliciano, J.D., and Siliciano, R.F. (2022). Measuring the latent reservoir for HIV-1: Quantification bias in near full-length genome sequencing methods. PLoS Pathog 18, e1010845.

25. Anderson, E.M., and Maldarelli, F. (2018). The role of integration and clonal expansion in HIV infection: live long and prosper. Retrovirology 15, 71.

26. Einkauf, K.B., Lee, G.Q., Gao, C., Sharaf, R., Sun, X., Hua, S., Chen, S.M., Jiang, C., Lian, X., Chowdhury, F.Z., et al. (2019). Intact HIV-1 proviruses accumulate at distinct chromosomal positions during prolonged antiretroviral therapy. J Clin Invest 129, 988–998.

27. Huang, A.S., Ramos, V., Oliveira, T.Y., Gaebler, C., Jankovic, M., Nussenzweig, M.C., and Cohn, L.B. (2021). Integration features of intact latent HIV-1 in CD4+ T cell clones contribute to viral persistence. J Exp Med 218.

28. Kuniholm, J., Armstrong, E., Bernabe, B., Coote, C., Berenson, A., Patalano, S.D., Olson, A., He, X., Lin, N.H., Fuxman Bass, J.I., and Henderson, A.J. (2021). Intragenic proviral elements support transcription of defective HIV-1 proviruses. PLoS Pathog 17, e1009982.

29. Collora, J.A., and Ho, Y.C. (2023). Integration site-dependent HIV-1 promoter activity shapes host chromatin conformation. Genome Res.

30. Vansant, G., Chen, H.C., Zorita, E., Trejbalová, K., Miklík, D., Filion, G., and Debyser, Z. (2020). The chromatin landscape at the HIV-1 provirus integration site determines viral expression. Nucleic Acids Res 48, 7801–7817.

31. Janssens, J., De Wit, F., Parveen, N., and Debyser, Z. (2022). Single-Cell Imaging Shows That the Transcriptional State of the HIV-1 Provirus and Its Reactivation Potential Depend on the Integration Site. mBio 13, e0000722.

32. Finzi, D., Hermankova, M., Pierson, T., Carruth, L.M., Buck, C., Chaisson, R.E., Quinn, T.C., Chadwick, K., Margolick, J., Brookmeyer, R., et al. (1997). Identification of a reservoir for HIV-1 in patients on highly active antiretroviral therapy. Science 278, 1295–1300.

33. Laird, G.M., Rosenbloom, D.I., Lai, J., Siliciano, R.F., and Siliciano, J.D. (2016). Measuring the Frequency of Latent HIV-1 in Resting CD4⁺ T Cells Using a Limiting Dilution Coculture Assay. Methods Mol Biol 1354, 239–253.

34. Enick, P.N., Brooker, J.P., Tumiotto, C.M., Staines, B.T., Eron, J.J., McMahon, D.K., Gandhi, R.T., Mellors, J.W., and Sobolewski, M.D. (2021). Comparison of methods to quantify inducible HIV-1 outgrowth. J Virus Erad 7, 100043.

35. Plantin, J., Massanella, M., and Chomont, N. (2018). Inducible HIV RNA transcription assays to measure HIV persistence: pros and cons of a compromise. Retrovirology 15, 9. 10.1186/s12977-017-0385-y.

36. Cillo, A.R., Sobolewski, M.D., Bosch, R.J., Fyne, E., Piatak Jr, M., Coffin, J.M., and Mellors, J.W. (2014). Quantification of HIV-1 latency reversal in resting CD4+ T cells from patients on suppressive antiretroviral therapy. Proceedings of the National Academy of Sciences 111, 7078–7083.

37. Massanella, M., Yek, C., Lada, S.M., Nakazawa, M., Shefa, N., Huang, K., and Richman, D.D. (2018). Improved assays to measure and characterize the inducible HIV reservoir. EBioMedicine 36, 113–121.

38. Procopio, F.A., Fromentin, R., Kulpa, D.A., Brehm, J.H., Bebin, A.-G., Strain, M.C., Richman, D.D., O’Doherty, U., Palmer, S., and Hecht, F.M. (2015). A novel assay to measure the magnitude of the inducible viral reservoir in HIV-infected individuals. EBioMedicine 2, 874–883.

39. Yucha, R.W., Hobbs, K.S., Hanhauser, E., Hogan, L.E., Nieves, W., Ozen, M.O., Inci, F., York, V., Gibson, E.A., and Thanh, C. (2017). High-throughput characterization of HIV-1 reservoir reactivation using a single-cell-in-droplet PCR assay. EBioMedicine 20, 217–229.

40. Bullen, C.K., Laird, G.M., Durand, C.M., Siliciano, J.D., and Siliciano, R.F. (2014). New ex vivo approaches distinguish effective and ineffective single agents for reversing HIV-1 latency in vivo. Nature medicine 20, 425–429.

41. Pasternak, A.O., Adema, K.W., Bakker, M., Jurriaans, S., Berkhout, B., Cornelissen, M., and Lukashov, V.V. (2008). Highly sensitive methods based on seminested real-time reverse transcription-PCR for quantitation of human immunodeficiency virus type 1 unspliced and multiply spliced RNA and proviral DNA. Journal of clinical microbiology 46, 2206–2211.

42. Shan, L., Rabi, S.A., Laird, G.M., Eisele, E.E., Zhang, H., Margolick, J.B., and Siliciano, R.F. (2013). A novel PCR assay for quantification of HIV-1 RNA. Journal of virology 87, 6521–6525.

43. Kaiser, P., Joshi, S.K., Kim, P., Li, P., Liu, H., Rice, A.P., Wong, J.K., and Yukl, S.A. (2017). Assays for precise quantification of total (including short) and elongated HIV-1 transcripts. Journal of virological methods 242, 1–8.

44. Lewin, S.R., Vesanen, M., Kostrikis, L., Hurley, A., Duran, M., Zhang, L., Ho, D.D., and Markowitz, M. (1999). Use of real-time PCR and molecular beacons to detect virus replication in human immunodeficiency virus type 1-infected individuals on prolonged effective antiretroviral therapy. Journal of virology 73, 6099–6103.

45. Zerbato, J.M., Khoury, G., Zhao, W., Gartner, M.J., Pascoe, R.D., Rhodes, A., Dantanarayana, A., Gooey, M., Anderson, J., Bacchetti, P., et al. (2021). Multiply spliced HIV RNA is a predictive measure of virus production ex vivo and in vivo following reversal of HIV latency. EBioMedicine 65, 103241.

46. Pasternak, A.O., Jurriaans, S., Bakker, M., Prins, J.M., Berkhout, B., and Lukashov, V.V. (2009). Cellular levels of HIV unspliced RNA from patients on combination antiretroviral therapy with undetectable plasma viremia predict the therapy outcome. PloS one 4, e8490.

47. Schmid, A., Gianella, S., von Wyl, V., Metzner, K.J., Scherrer, A.U., Niederöst, B., Althaus, C.F., Rieder, P., Grube, C., and Joos, B. (2010). Profound depletion of HIV-1 transcription in patients initiating antiretroviral therapy during acute infection. PloS one 5, e13310.

48. Lungu, C., and Procopio, F.A. (2022). TILDA: Tat/Rev Induced Limiting Dilution Assay. Methods Mol Biol 2407, 365–372.

49. Baxter, A.E., Niessl, J., Fromentin, R., Richard, J., Porichis, F., Massanella, M., Brassard, N., Alsahafi, N., Routy, J.P., Finzi, A., et al. (2017). Multiparametric characterization of rare HIV-infected cells using an RNA-flow FISH technique. Nat Protoc 12, 2029–2049.

50. Baxter, A.E., Niessl, J., Morou, A., and Kaufmann, D.E. (2017). RNA flow cytometric FISH for investigations into HIV immunology, vaccination and cure strategies. AIDS Res Ther 14, 40.

51. Sannier, G., Dubé, M., Dufour, C., Richard, C., Brassard, N., Delgado, G.G., Pagliuzza, A., Baxter, A.E., Niessl, J., Brunet-Ratnasingham, E., et al. (2021). Combined single-cell transcriptional, translational, and genomic profiling reveals HIV-1 reservoir diversity. Cell Rep 36, 109643.

52. Abdel-Mohsen, M., Richman, D., Siliciano, R.F., Nussenzweig, M.C., Howell, B.J., Martinez-Picado, J., Chomont, N., Bar, K.J., Yu, X.G., Lichterfeld, M., et al. (2020). Recommendations for measuring HIV reservoir size in cure-directed clinical trials. Nat Med 26, 1339–1350.

53. Rossouw, T., Tucker, J.D., van Zyl, G.U., Sikwesi, K., and Godfrey, C. (2017). Barriers to HIV remission research in low- and middle-income countries. J Int AIDS Soc 20, 21521.

54. Mori, Y., and Notomi, T. (2009). Loop-mediated isothermal amplification (LAMP): a rapid, accurate, and cost-effective diagnostic method for infectious diseases. J Infect Chemother 15, 62–69.

55. Dao Thi, V.L., Herbst, K., Boerner, K., Meurer, M., Kremer, L.P., Kirrmaier, D., Freistaedter, A., Papagiannidis, D., Galmozzi, C., Stanifer, M.L., et al. (2020). A colorimetric RT-LAMP assay and LAMP-sequencing for detecting SARS-CoV-2 RNA in clinical samples. Sci Transl Med 12.

56. Yan, C., Cui, J., Huang, L., Du, B., Chen, L., Xue, G., Li, S., Zhang, W., Zhao, L., Sun, Y., et al. (2020). Rapid and visual detection of 2019 novel coronavirus (SARS-CoV-2) by a reverse transcription loop-mediated isothermal amplification assay. Clin Microbiol Infect 26, 773–779.

57. Huang, W.E., Lim, B., Hsu, C.C., Xiong, D., Wu, W., Yu, Y., Jia, H., Wang, Y., Zeng, Y., Ji, M., et al. (2020). RT-LAMP for rapid diagnosis of coronavirus SARS-CoV-2. Microb Biotechnol 13, 950–961.

58. Joung, J., Ladha, A., Saito, M., Kim, N.-G., Woolley, A.E., Segel, M., Barretto, R.P.J., Ranu, A., Macrae, R.K., Faure, G., et al. (2020). Detection of SARS-CoV-2 with SHERLOCK One-Pot Testing. New England Journal of Medicine 383, 1492–1494. 10.1056/NEJMc2026172.

59. Amoah, I.D., Mthethwa, N.P., Pillay, L., Deepnarain, N., Pillay, K., Awolusi, O.O., Kumari, S., and Bux, F. (2021). RT-LAMP: A Cheaper, Simpler and Faster Alternative for the Detection of SARS-CoV-2 in Wastewater. Food Environ Virol 13, 447–456.

60. Rudolph, D.L., Sullivan, V., Owen, S.M., and Curtis, K.A. (2015). Detection of Acute HIV-1 Infection by RT-LAMP. PLoS One 10, e0126609.

61. Curtis, K.A., Morrison, D., Rudolph, D.L., Shankar, A., Bloomfield, L.S.P., Switzer, W.M., and Owen, S.M. (2018). A multiplexed RT-LAMP assay for detection of group M HIV-1 in plasma or whole blood. J Virol Methods 255, 91–97.

62. Li, Y., Chen, X., Zhao, Y., Wan, Z., Zeng, Y., Ma, Y., Zhou, L., Xu, G., Reboud, J., Cooper, J.M., and Zhang, C. (2021). A rapid variant-tolerant reverse transcription loop-mediated isothermal amplification assay for the point of care detection of HIV-1. Analyst 146, 5347–5356. 10.1039/d1an00598g.

63. Bertoldi, A., D’Urbano, V., Bon, I., Verbon, A., Rokx, C., Boucher, C., van Kampen, J.J.A., Gruters, R.A., Gallinella, G., Calza, L., et al. (2020). Development of C-TILDA: A modified TILDA method for reservoir quantification in long term treated patients infected with subtype C HIV-1. Journal of Virological Methods 276, 113778. https://doi.org/10.1016/j.jviromet.2019.113778.

64. Lungu, C., Banga, R., Gruters, R.A., and Procopio, F.A. (2021). Inducible HIV-1 Reservoir Quantification: Clinical Relevance, Applications and Advancements of TILDA. Front Microbiol 12, 686690.

65. Lungu, C., Procopio, F.A., Overmars, R.J., Beerkens, R.J.J., Voermans, J.J.C., Rao, S., Prins, H.A.B., Rokx, C., Pantaleo, G., Vijver, D., et al. (2020). Inter-Laboratory Reproducibility of Inducible HIV-1 Reservoir Quantification by TILDA. Viruses 12.

66. Prins, H.A.B., Crespo, R., Lungu, C., Rao, S., Li, L., Overmars, R.J., Papageorgiou, G., Mueller, Y.M., Stoszko, M., Hossain, T., et al. (2023). The BAF complex inhibitor pyrimethamine reverses HIV-1 latency in people with HIV-1 on antiretroviral therapy. Sci Adv 9, eade6675.

67. Gadkar, V.J., Goldfarb, D.M., Gantt, S., and Tilley, P.A.G. (2018). Real-time Detection and Monitoring of Loop Mediated Amplification (LAMP) Reaction Using Self-quenching and De-quenching Fluorogenic Probes. Scientific Reports 8, 5548. 10.1038/s41598-018-23930-1.

68. Kim, J., Park, B.G., Lim, D.H., Jang, W.S., Nam, J., Mihn, D.C., and Lim, C.S. (2021). Development and evaluation of a multiplex loop-mediated isothermal amplification (LAMP) assay for differentiation of Mycobacterium tuberculosis and non-tuberculosis mycobacterium in clinical samples. PLoS One 16, e0244753.

69. Sherrill-Mix, S., Hwang, Y., Roche, A.M., Glascock, A., Weiss, S.R., Li, Y., Haddad, L., Deraska, P., Monahan, C., Kromer, A., et al. (2021). Detection of SARS-CoV-2 RNA using RT-LAMP and molecular beacons. Genome Biol 22, 169.

70. Tanner, N.A., Zhang, Y., and Evans, T.C., Jr. (2012). Simultaneous multiple target detection in real-time loop-mediated isothermal amplification. Biotechniques 53, 81–89.

71. Gandhi, R.T., McMahon, D.K., Bosch, R.J., Lalama, C.M., Cyktor, J.C., Macatangay, B.J., Rinaldo, C.R., Riddler, S.A., Hogg, E., Godfrey, C., et al. (2017). Levels of HIV-1 persistence on antiretroviral therapy are not associated with markers of inflammation or activation. PLOS Pathogens 13, e1006285.

72. Scully, E.P., Gandhi, M., Johnston, R., Hoh, R., Lockhart, A., Dobrowolski, C., Pagliuzza, A., Milush, J.M., Baker, C.A., Girling, V., et al. (2018). Sex-Based Differences in Human Immunodeficiency Virus Type 1 Reservoir Activity and Residual Immune Activation. The Journal of Infectious Diseases 219, 1084–1094. 10.1093/infdis/jiy617.

73. Falcinelli, S.D., Shook-Sa, B.E., Dewey, M.G., Sridhar, S., Read, J., Kirchherr, J., James, K.S., Allard, B., Ghofrani, S., Stuelke, E., et al. (2020). Impact of Biological Sex on Immune Activation and Frequency of the Latent HIV Reservoir During Suppressive Antiretroviral Therapy. J Infect Dis 222, 1843–1852.

74. Prodger, J.L., Capoferri, A.A., Yu, K., Lai, J., Reynolds, S.J., Kasule, J., Kityamuweesi, T., Buule, P., Serwadda, D., Kwon, K.J., et al. (2020). Reduced HIV-1 latent reservoir outgrowth and distinct immune correlates among women in Rakai, Uganda. JCI Insight 5.

75. Siliciano, J.D., and Siliciano, R.F. (2021). Low Inducibility of Latent Human Immunodeficiency Virus Type 1 Proviruses as a Major Barrier to Cure. J Infect Dis 223, 13–21.

76. Das, B., Dobrowolski, C., Luttge, B., Valadkhan, S., Chomont, N., Johnston, R., Bacchetti, P., Hoh, R., Gandhi, M., Deeks, S.G., et al. (2018). Estrogen receptor-1 is a key regulator of HIV-1 latency that imparts gender-specific restrictions on the latent reservoir. Proc Natl Acad Sci U S A 115, E7795–E7804.

77. Szotek, E.L., Narasipura, S.D., and Al-Harthi, L. (2013). 17β-Estradiol inhibits HIV-1 by inducing a complex formation between β-catenin and estrogen receptor α on the HIV promoter to suppress HIV transcription. Virology 443, 375–383.

78. Ndung’u, T., McCune, J.M., and Deeks, S.G. (2019). Why and where an HIV cure is needed and how it might be achieved. Nature 576, 397–405.

79. Lifson, M.A., Ozen, M.O., Inci, F., Wang, S., Inan, H., Baday, M., Henrich, T.J., and Demirci, U. (2016). Advances in biosensing strategies for HIV-1 detection, diagnosis, and therapeutic monitoring. Adv Drug Deliv Rev 103, 90–104.

80. Tamura, K., Stecher, G., and Kumar, S. (2021). MEGA11: Molecular Evolutionary Genetics Analysis Version 11. Molecular Biology and Evolution 38, 3022–3027. 10.1093/molbev/msab120.

81. Jordan, A., Bisgrove, D., and Verdin, E. (2003). HIV reproducibly establishes a latent infection after acute infection of T cells in vitro. Embo J 22, 1868–1877.

82. Rosenbloom, D.I.S., Elliott, O., Hill, A.L., Henrich, T.J., Siliciano, J.M., and Siliciano, R.F. (2015). Designing and Interpreting Limiting Dilution Assays: General Principles and Applications to the Latent Reservoir for Human Immunodeficiency Virus-1. Open Forum Infectious Diseases 2. 10.1093/ofid/ofv123.

