## Supplementary figures and images for "SQuHIVLa: A novel assay for Specific Quantification of inducible HIV-1 reservoir by LAMP"

### Supplemental Figure S1

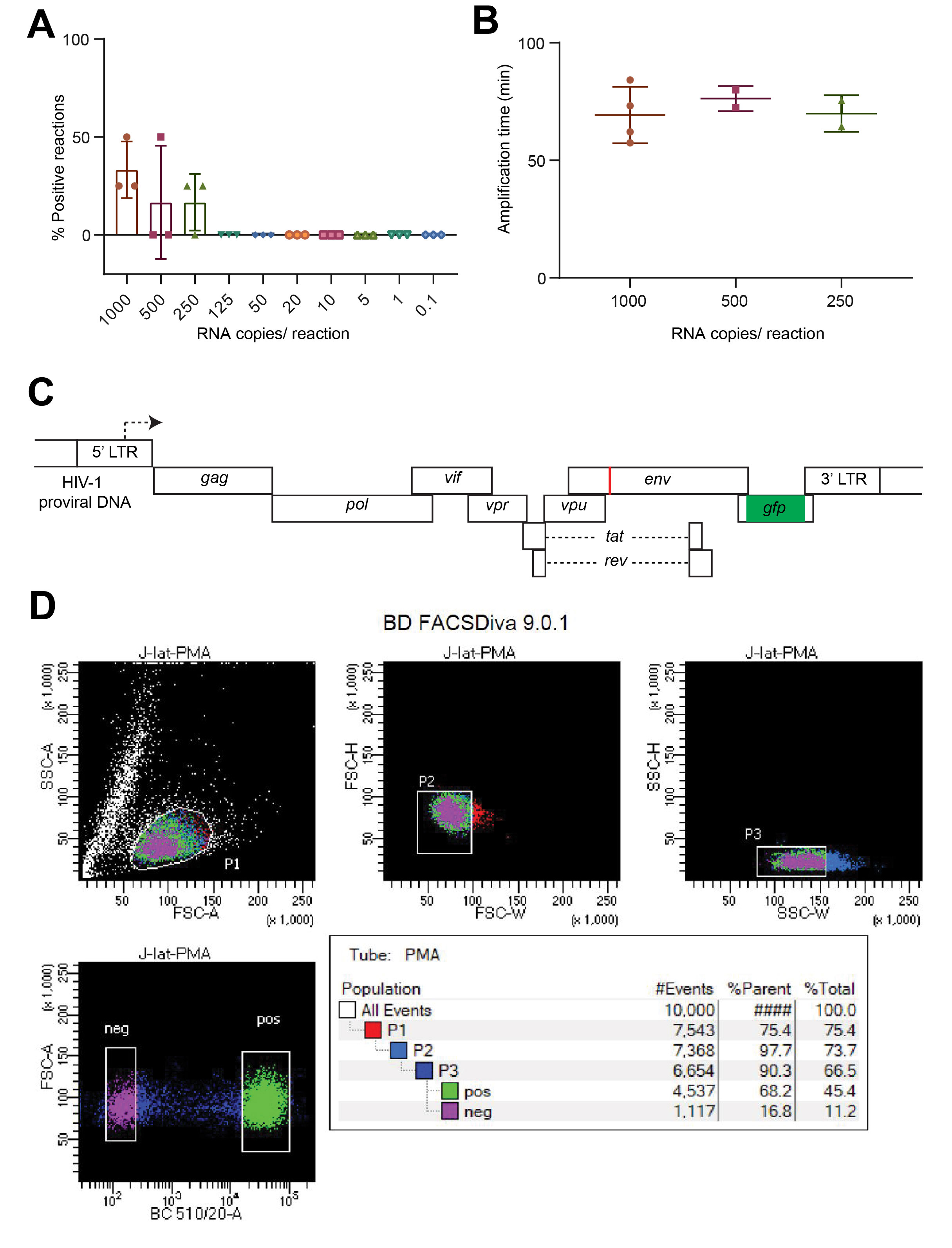

### Supplemental Figure S2

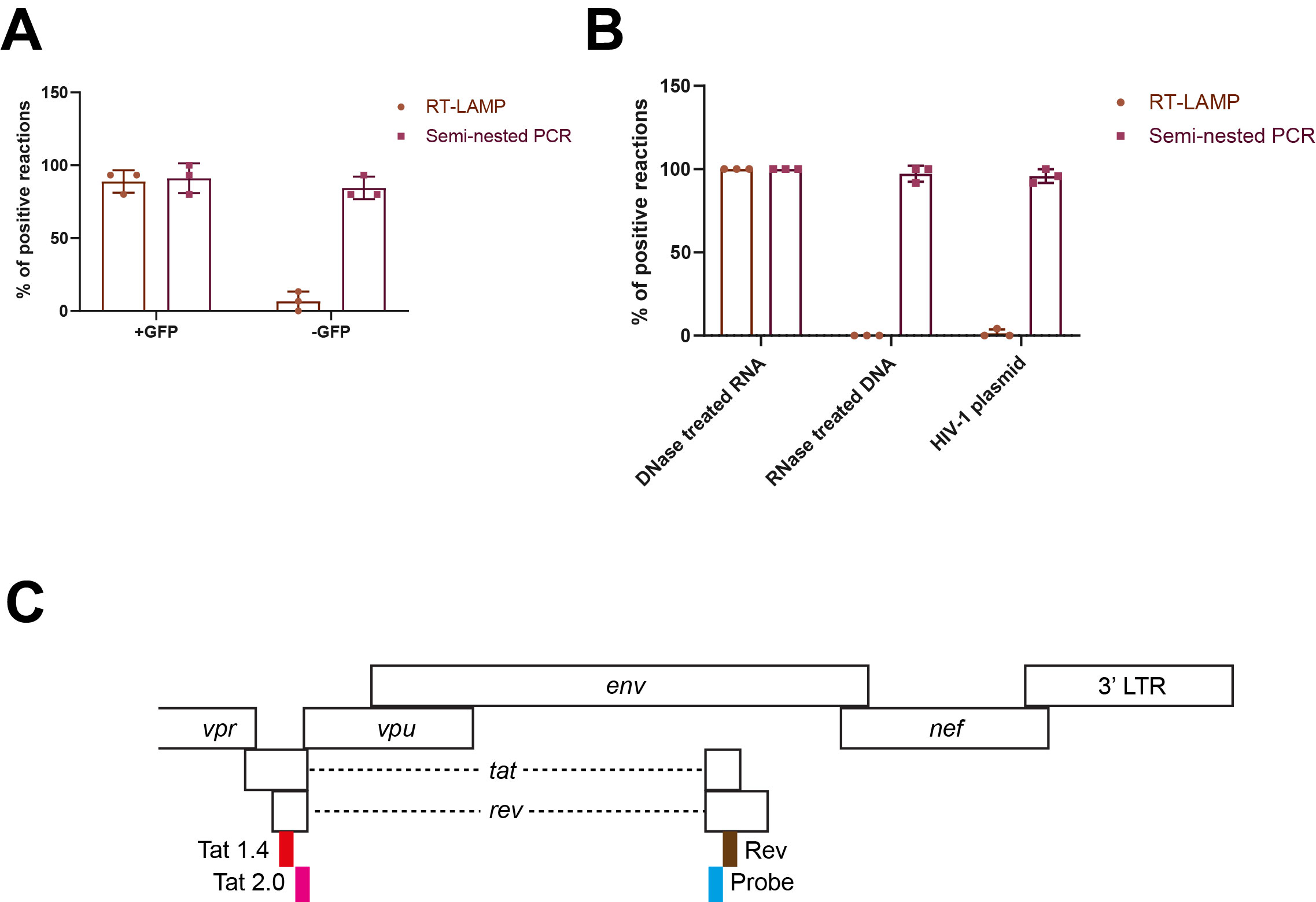

### Supplemental Figure S3

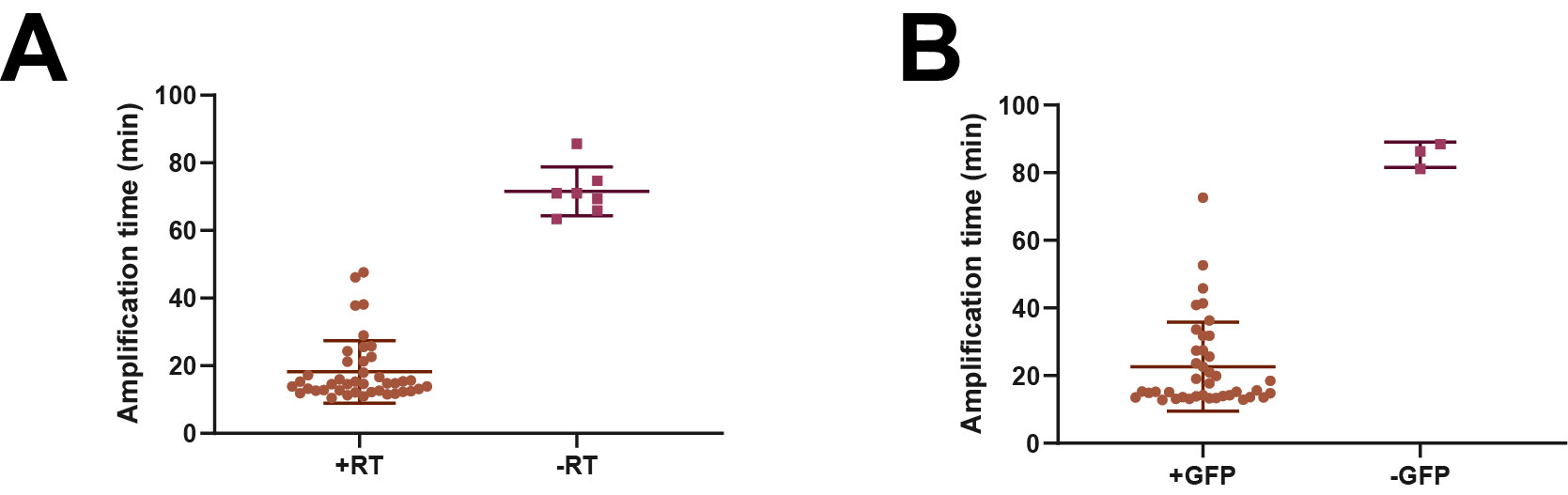

### Supplemental Figure S4

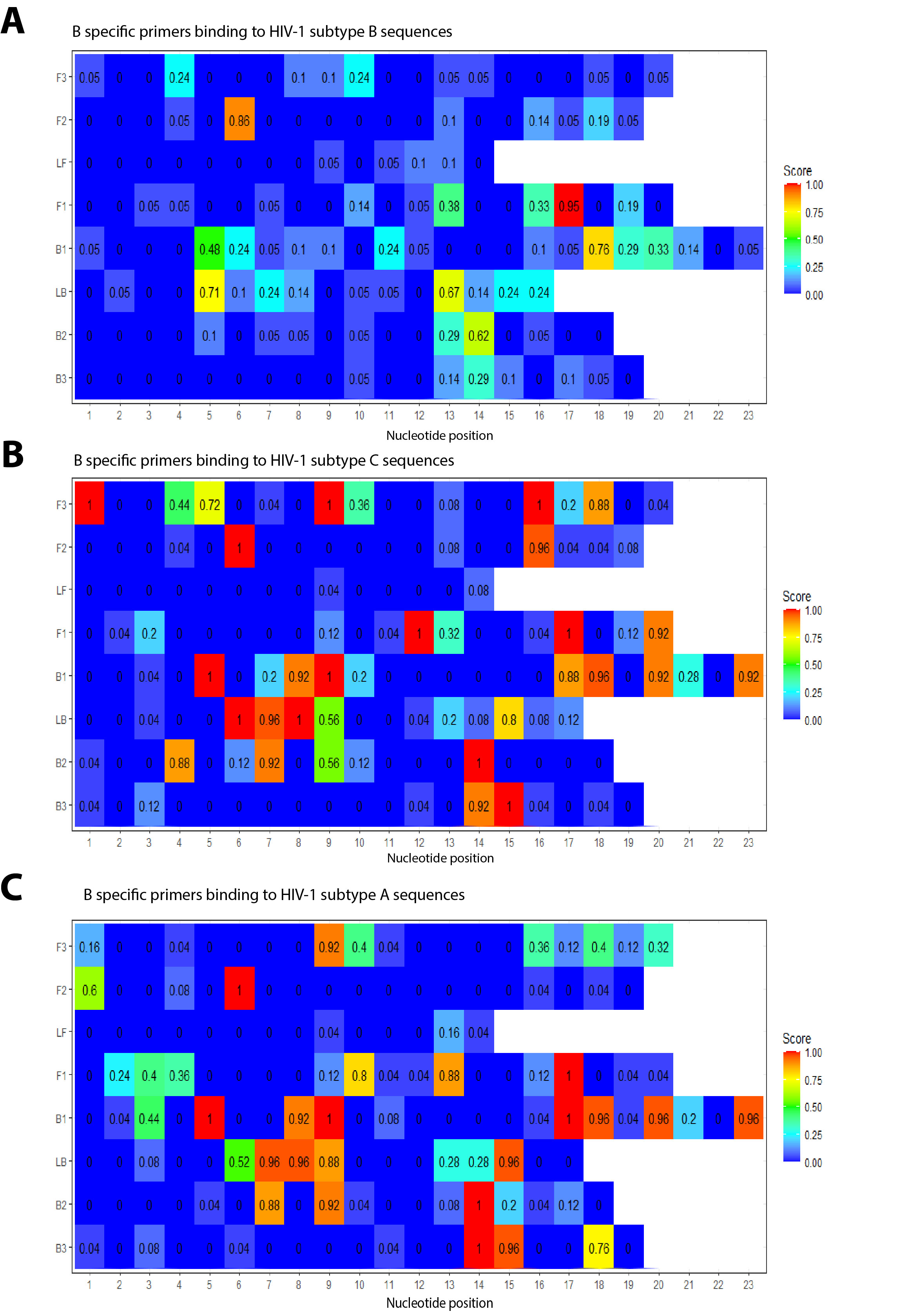

### Supplemental Figure S5

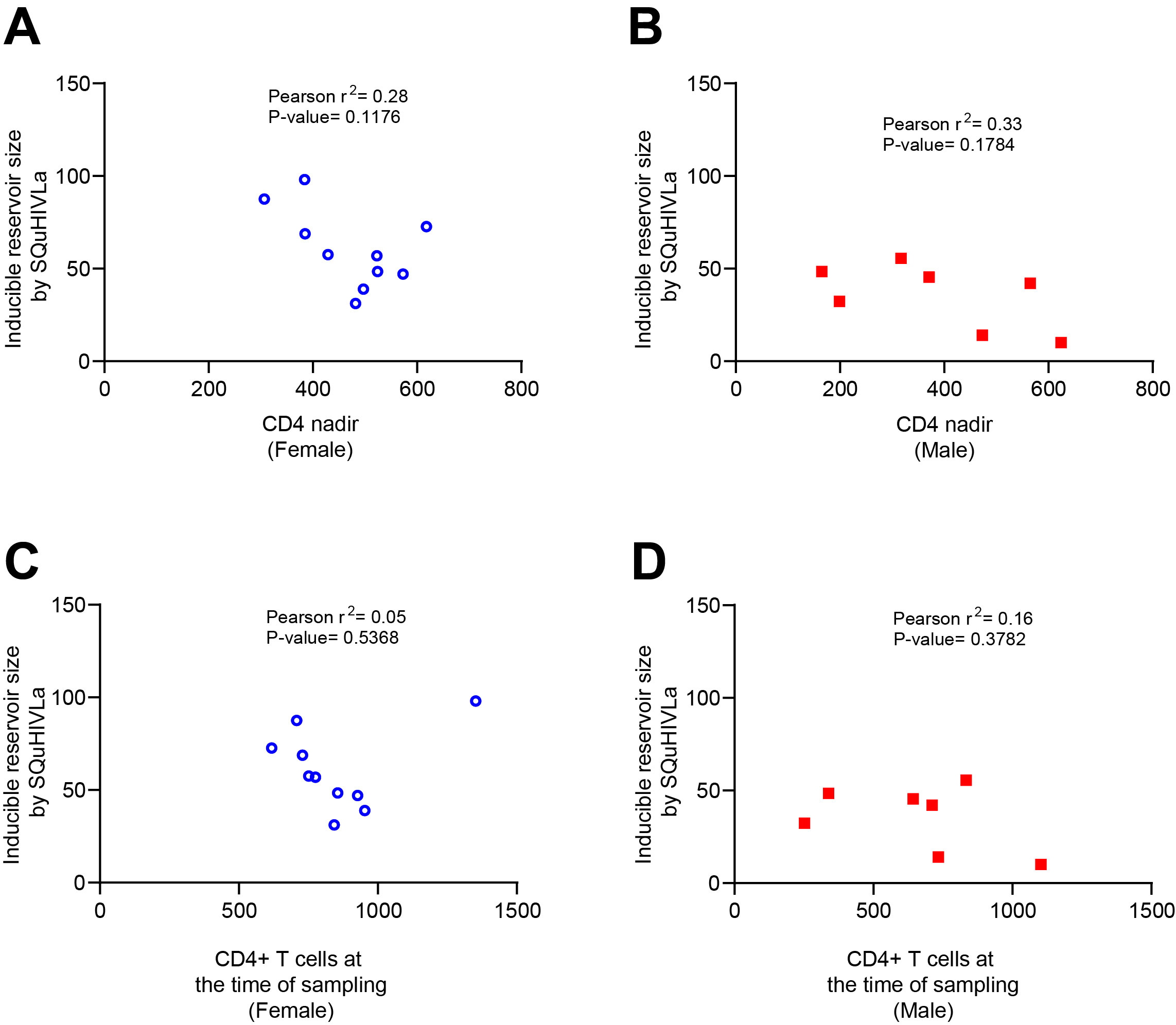
